# GCAD: a Computational Framework for Mammalian Genetic Program Computer-Aided Design

**DOI:** 10.1101/2025.08.23.671908

**Authors:** Kathleen S. Dreyer, Anh V. Nguyen, Gauri G. Bora, Lauren E. Redus, Hailey I. Edelstein, Jocelyn J. Garcia, Eleftheria Anastasia, Kate E. Dray, Joshua N. Leonard, Niall M. Mangan

## Abstract

Genetic programs can direct living systems to perform diverse, pre-specified functions. As the library of parts available for building such programs continues to expand, computation-guided design is increasingly helpful and necessary. Predictive models aid the challenging design process, but iterative simulation and experimentation are intractable for complex functions. Computer-aided design accelerates this process, but existing tools do not yet capture the behavior of mammalian-specific parts and population-level effects needed for mammalian synthetic biologists. To address these needs, we developed a framework for mammalian genetic program computer-aided design. Starting with a user-defined design specification to quantify circuit performance, the framework uses a genetic algorithm to search through possible designs. Circuit space is defined by a library of experimentally characterized parts and dynamical systems models for gene expression in a heterogeneous cell population. We developed this genetic algorithm using a directed graph-based formulation with biologically constrained rules to explore regulatory connections and parts. We evaluated the framework for design problems of varying complexity, including programs we describe as an amplifier, signal conditioner, and pulse generator, demonstrating that the algorithm can successfully find optimal circuit designs. Finally, we experimentally evaluated selected circuits, demonstrating the path from a predicted circuit design to experimental testing and highlighting the importance of characterization for enabling predictive design. Overall, this framework establishes general approaches that can be refined and expanded, accelerating the design and implementation of mammalian genetic programs.

## Introduction

Genetic programs (genetic circuits) are groups of biological parts—including DNA, RNA, and regulatory proteins—that enable control of customizable cellular functions, and the utility of this concept is well-established in synthetic biology.^1^ Nonetheless, the process of designing these programs remains challenging, particularly in mammalian systems. The design, build, test, learn (DBTL) cycle is employed to build genetic programs and understand their function, yet iteration is rate limiting and laborious in mammalian systems.^2, 3^ One factor that commonly necessitates iteration is discrepancies between expected and observed behavior of genetic circuits driven by phenomena not considered explicitly at the design phase, such as cell population-level heterogeneity (e.g., variation in DNA plasmid uptake across a population of cells in transfection-based experiments).^2, 4^ Moreover, the experimental DBTL cycle becomes intractable when considering large design spaces (i.e., considering many parts and topologies through which they may be connected) or when optimizing for complex desired functions that may involve multiple design objectives, including desired dynamic behaviors.^3^

Model-guided, predictive design can accelerate the experimental DBTL cycle of genetic programs but is limited by a researcher’s ability to propose program designs. Mathematical models are useful tools to gain insight into mechanisms underlying system function and to propose designs that have not been experimentally evaluated.^5^ Ordinary differential equations (ODEs) are well-suited for describing the dynamics of genetic programs, since ODEs describe the time dependent evolution of system components based on a proposed set of mechanisms. Predictive design of mammalian genetic programs has previously been demonstrated,^6–11^ in each case reducing the amount of experimental trial-and-error and highlighting the promise of this approach. Models that incorporate descriptions of cell population-level heterogeneity^6, 11^ are particularly useful for improving understanding of the often-unintuitive impact of this factor on program performance and have inspired designs that mitigate such effects. Despite these advantages, model-guided genetic program design is often limited by the intuition of the researcher—one must propose circuit topologies and/or theoretical parts to evaluate in simulation, and iteration is often required to achieve a circuit design with a predicted performance that meets the design goal, particularly for complex desired functions. A potential alternative to using a small number of human-designed circuits is an exhaustive combinatorial (hereafter, “combinatorial”) search of the genetic program design space, in which all possible combinations of parts and component doses in all feasible circuit topologies are systematically evaluated. However, a combinatorial search becomes computationally intractable for design spaces consisting of several parts and potential regulatory interactions, easily reaching millions of potential circuits. Design automation through a computationally assisted search or optimization expands predictive capability beyond intuition alone while avoiding a combinatorial search; several approaches have been developed for this purpose, but none meet the need of mammalian cellular engineers.

The existing literature on automated design of genetic circuits can be grouped into two categories, ideal component characterization and functional topology search. The former aims to design an ideal circuit that satisfies user inputs by optimizing the kinetic parameters of the genetic components in the circuit.^12–17^ Specifically, these methods often determine the circuit topology in advance, i.e., the number of component types (e.g., promoters, activators, or repressors) and how they are connected in the circuit are known. However, the corresponding kinetic parameters of each component such as basal production, degradation rate, or cooperativity coefficient are variables to be optimized. While ideal component characterization can be useful to predict functionality that cannot be achieved with existing parts, it requires additional experimental design to develop parts that align with the predicted, theoretical parts, which often involves time-intensive, trial-and-error iterations.

Unlike the first category, functional topology searches start with a library of available, characterized components and attempt to find a combination of components and their connections that meet the design specification. While methods for component characterization are predominantly built on gradient-based optimization, the methods for topology search utilize a wider range of techniques. For instance, Huynh et al. formulated an integer nonlinear optimization problem with binary variables representing the presence/absence of a component or a connection in the circuit.^18^ Heuristic searches like simulated annealing,^19–21^ directed evolution,^22–25^ and genetic algorithm^26, 27^ are also popular. Simulated annealing is an optimization technique which at every iteration perturbs the previous solution candidate, calculates the objective, and accepts the new candidate with a probability proportional to the objective. Directed evolution is an ensemble method that evolves a population of candidate solutions by generating random mutations and exploring the search space. Genetic algorithm (GA) is similar to directed evolution but includes a crossover operator, which promotes exploitation of good candidates. Both GA and directed evolution select top performing candidates to advance to the next iteration, or generation, as referred to in GA literature. Prior work using the functional topology search approach generally employed either existing models of foundational synthetic bacterial circuits,^19, 21, 28^ or models of bacterial or yeast logic gates,^20, 29^ neither of which are readily applicable to mammalian systems. Additionally, several of these tools use Boolean logic functions as design specifications,^19–21^ but it is often of interest in mammalian synthetic biology to design circuits with dynamic behaviors, e.g., for applications in stem cell differentiation.^30^

It should be noted that topology search and component characterization are not mutually exclusive. For example, Hiscock proposed a method that starts with the most generalized circuit comprising of all available components and used an ℓ_l_ regularized optimization to shrink kinetic coefficients and remove weak connections.^31^ This effectively sparsified the original circuit and output a simplified topology with fewer connections and potentially fewer components. While parameter regularization makes the optimization more robust, topology sparsification may not be guaranteed. Shen et al. took a more direct approach such that for each iteration, their method found the connection whose removal causes minimal change in the circuit’s performance by taking out one connection at a time.^32^ The method then retrained kinetic parameters of the new circuit. This led to a sequence of topologies with a decreasing number of connections and, thus, guaranteed a simplified outcome. However, the repeated leave-one-out connection test can become computationally expensive. Dasika et al. solved a mixed integer dynamic optimization problem to get a list of circuits corresponding to local minima.^33^ The method also included a protocol for fine-tuning kinetic parameters such as promoter strength. Marchisio et al. generated a set of potential candidates from known parts and performed sensitivity analysis and parameter optimization to further fine-tune the solution.^34^

An ideal framework for mammalian genetic program design would utilize a library of parts experimentally characterized in mammalian cells to limit the need for trial-and-error tuning of theoretical parts and would be amenable to dynamic and steady-state design specifications. To address this need, we developed a framework for mammalian Genetic program Computer-Aided Design (GCAD). We utilize a functional circuit topology search with a library of previously developed, synthetic transcription factor (synTF) activators and repressors for mammalian cells (referred to as synTF-As and synTF-Rs, respectively) and an associated genetic parts model.^35^ This choice of experimentally characterized parts is well suited to developing GCAD as it comprises a large set of components that are functionally similar, but which differ in quantitative parameters associated with each component. Such a set enables experimental implementation of GCAD-predicted circuit designs, rather than necessitating experimental design and characterization of novel parts with functionality that aligns with that of the mathematically modeled parts. Although the model associated with this toolkit was focused on explanatory goals^35^ and did not employ a formal parameter identifiability analysis,^5^ this nonetheless serves as a useful base case for driving GCAD development.

We considered design goals for steady-state and dynamic behavior in the development of GCAD, ensuring that the types of design goals relevant in mammalian synthetic biology can be utilized for circuit design with this framework. We developed a graph-based GA circuit selection method to search the circuit design space and identify optimal designs. Graph-based GAs have been developed in the context of automated design of chemical molecules using undirected graphs, and here we developed a method for directed graphs with additional edge properties (See **Supplementary Note 1** for discussion of existing methods). Using a GA offers a few key advantages over other approaches. In particular, the GA can be flexibly adapted for different types of optimization objectives because the method is not gradient-based. Furthermore, it is straightforward to extend GAs to enable multi-objective optimization via sorting methods such as Non-dominated Sorting GA (NSGA) II,^36^ which expands the types of design goals enabled with the method. To our knowledge, our work is the first to formally encode circuit topologies as directed graphs and appropriately modify the GA operators based on this encoding. This GA methodology comprises a unique contribution to the field of genetic program design automation, exploring a promising GA search strategy for systems with a natural graph representation.

We assessed the GCAD framework for 3 test cases of varying complexity, each of which is motivated by a relevant design challenge in mammalian synthetic biology. We evaluated the performance of the GA with a simplified version of the aforementioned genetic parts model and compared to a combinatorial search to gain confidence that the GA can recover optimal circuit designs. We then incorporated a representation of cell population-level heterogeneity to design circuits based on average population-level behavior. For test cases in which the most optimal topologies found meet the design goal, we demonstrated that selected designs can be built and experimentally evaluated. Taken together, the GCAD framework meets key needs of mammalian synthetic biologists and is a first step towards a generalizable tool for automated mammalian genetic program design.

## Results

### Framing the circuit design problem mathematically within the GCAD framework

The GCAD framework takes a design objective and a library of mammalian genetic parts as inputs and searches across circuit topologies and parts using a genetic algorithm to find potential optimal designs; the user then evaluates candidate designs manually (i.e., to explore features not explicitly framed as constraints, but which may make a design attractive or not) and implements suitable designs experimentally (**Figure 1**). The first input to GCAD is the Design Specification, where the desired qualitative function is translated into a quantitative objective (or objectives) for optimization. For example, if one wishes to amplify the magnitude of expression of a fluorescent reporter gene driven by a specified promoter when it is activated (the ON-state), we quantified this objective using the ratio of the reporter gene expression of an amplification circuit in the ON_state to the reporter expression of a reference case without amplification in the ON_state (ON_rel_). The objective function provided as input to the circuit selection method is then to maximize ON_rel_.

**Figure 1.**
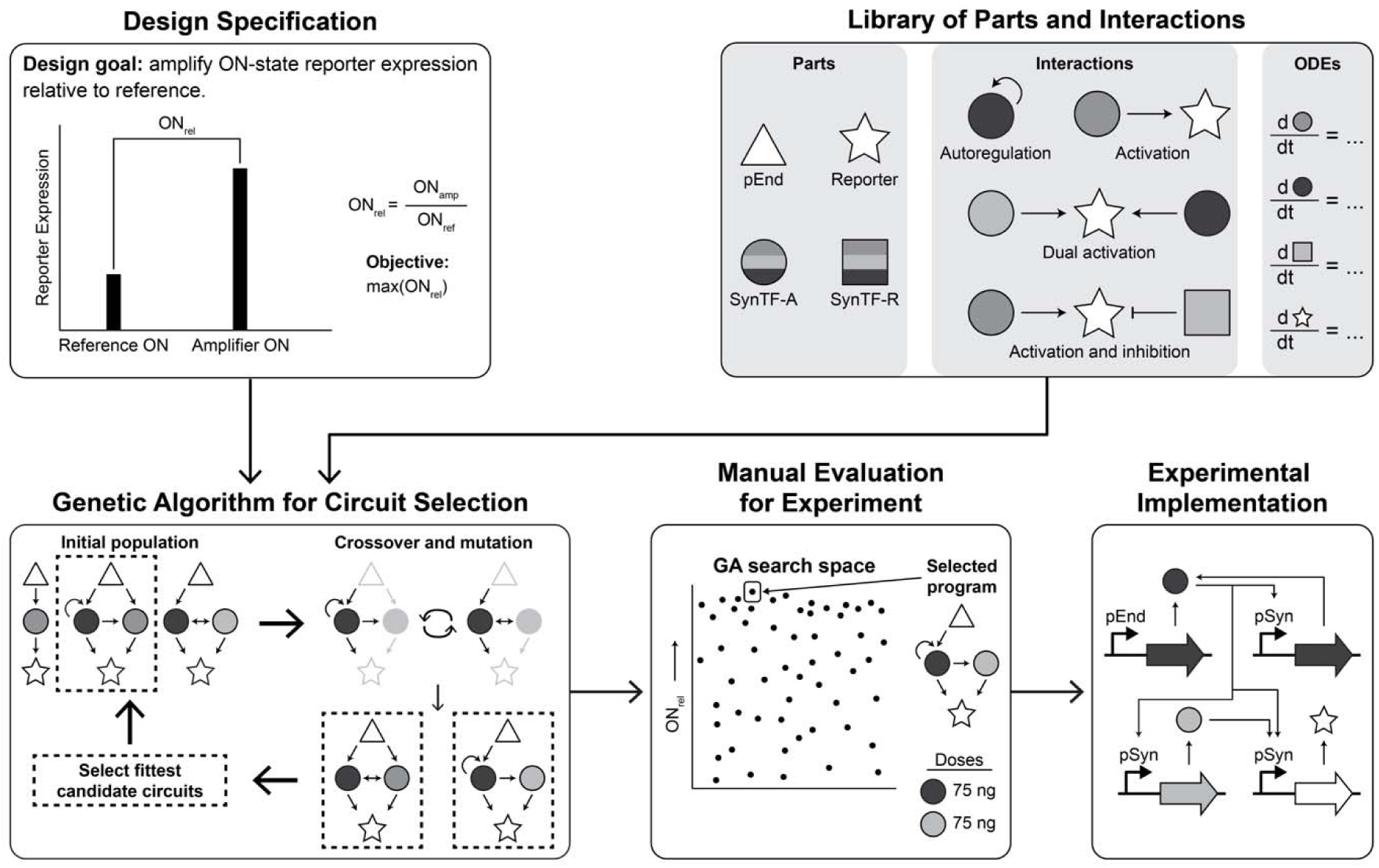
GCAD provides a framework for specifying and guiding genetic program design. The GCAD (Genetic program Computer-Aided Design) framework utilizes a design specification and a library of parts and potential interactions (described by an ODE model) as inputs to a genetic algorithm that searches through circuit designs. The range of circuit candidates explored during the search **is** manually evaluated and a subset of optimal circuits are selected for experimental implementation. The selected genetic circuits are constructed as DNA plasmids for experimental testing. GA: genetic algorithm; pEnd: endogenous promoter; pSyn: synthetic promoter; synTF-A: synthetic transcription factor activator; synTF-R: synthetic transcription factor repressor.

The second input to GCAD is the library of genetic parts and rules for regulatory interactions, which are represented mathematically using a genetic parts model for a suite of transcriptional activators and repressors, and a reporter protein (**Figure 1**). A genetic program can be represented as a system of ordinary differential equations based on an existing, genetic parts model (**Methods**). We included representations of parts and interactions previously characterized^35^, and we also extended the model to describe how synTF-As and/or synTF-Rs might co-regulate a jointly targeted promoter; to this end we incorporated hypothesized representations of (i) dual activator or (ii) activator and repressor regulatory interactions comprising different synTF variants. Although such interactions were not characterized in the initial study (excepting a few special cases involving synTF1 and synTF2), we opted to include these speculative extensions to create a full design space for developing GCAD. Therefore, our expectation was that the solutions from the GA would comprise behaviors that could be achieved if the parts functioned *as hypothesized*. We developed a set of rules to automatically generate model equations based on the characteristics of a given circuit, which is used in the circuit selection method to translate circuit designs in the search space into ODEs to simulate their performance.

Our GA-based method searches the circuit design space to select a program with the optimal objective—or the programs with Pareto optimal objectives, in the case of multi-objective optimization (**Methods**). A GA is a more computationally efficient alternative to a combinatorial search, which becomes intractable for complex design spaces and/or if the simulation time of each circuit is too high. The GA starts with an initial population and iteratively updates this population by generating and testing new circuits through mutation and combination of the best performing circuits at each generation. To enable further manual evaluation of the search space explored by GCAD, we output the full search space of programs and corresponding objectives, so that experimentalist can select a set of programs with different objective function values in addition to that of the optimal solution(s) found. The selected genetic program(s) can then be translated into a set of DNA constructs for experimental implementation (**Figure 1**).

### Test case 1: amplification of output gene expression

To demonstrate and evaluate the GCAD framework, we developed a set of test cases that represent design challenges in mammalian synthetic biology of varying complexity, including amplification (i.e., to make output of an existing module greater), and inducible dynamic behavior (in this case a pulse in gene expression of limited duration). For the test cases, we considered a scenario in which a researcher is investigating an endogenous promoter and its regulation of the expression of an endogenous gene. The researcher may track the activity of the endogenous promoter by placing the expression of a fluorescent reporter under its control; we refer to this system as the reference case (i.e., a standard fluorescent reporter). For simplicity, we consider the promoter to exist in either in an OFF-state or an ON-state (rather than considering the dynamics of the state transition). Common challenges that one may encounter in practice are that the fluorescent reporter output may be insufficient to distinguish the ON-state from the OFF-state, or the ON-state may be too modest to quantify changes (e.g., dynamics) of interest. In these cases, the researcher may benefit from a genetic program to process information from the endogenous promoter and transform it into a more useful output. A researcher could typically define specific design goals to address the needs of their investigation, using the reference case as a benchmark for performance. This framing forms the basis for the following test cases.

We first defined a simple design goal for an information processing gene circuit—amplification of reporter expression in the ON-state relative to the reference case; we refer to this test case as the Amplifier. For the design specification, we defined the ratio of the Amplifier reporter expression in the ON-state to that of the reference (ON_rel_) as a quantitative metric for the design goal, with the objective being to select a genetic program with the maximum ON_rel_ value— circuits with ON_rel_ greater than 1 would thereby satisfy the design goal (**Figure 2a**). We considered a design space consisting of the activators and repressors (12 synTF-As and 12 synTF-Rs, respectively), 156 promoters that can be regulated by any pair of synTFs (or by a single synTF binding to all sites), and all possible doses and interactions of the synTF-As and synTF-Rs. We assumed no prior intuition as to which parts are likely to meet the design specification.

**Figure 2.**
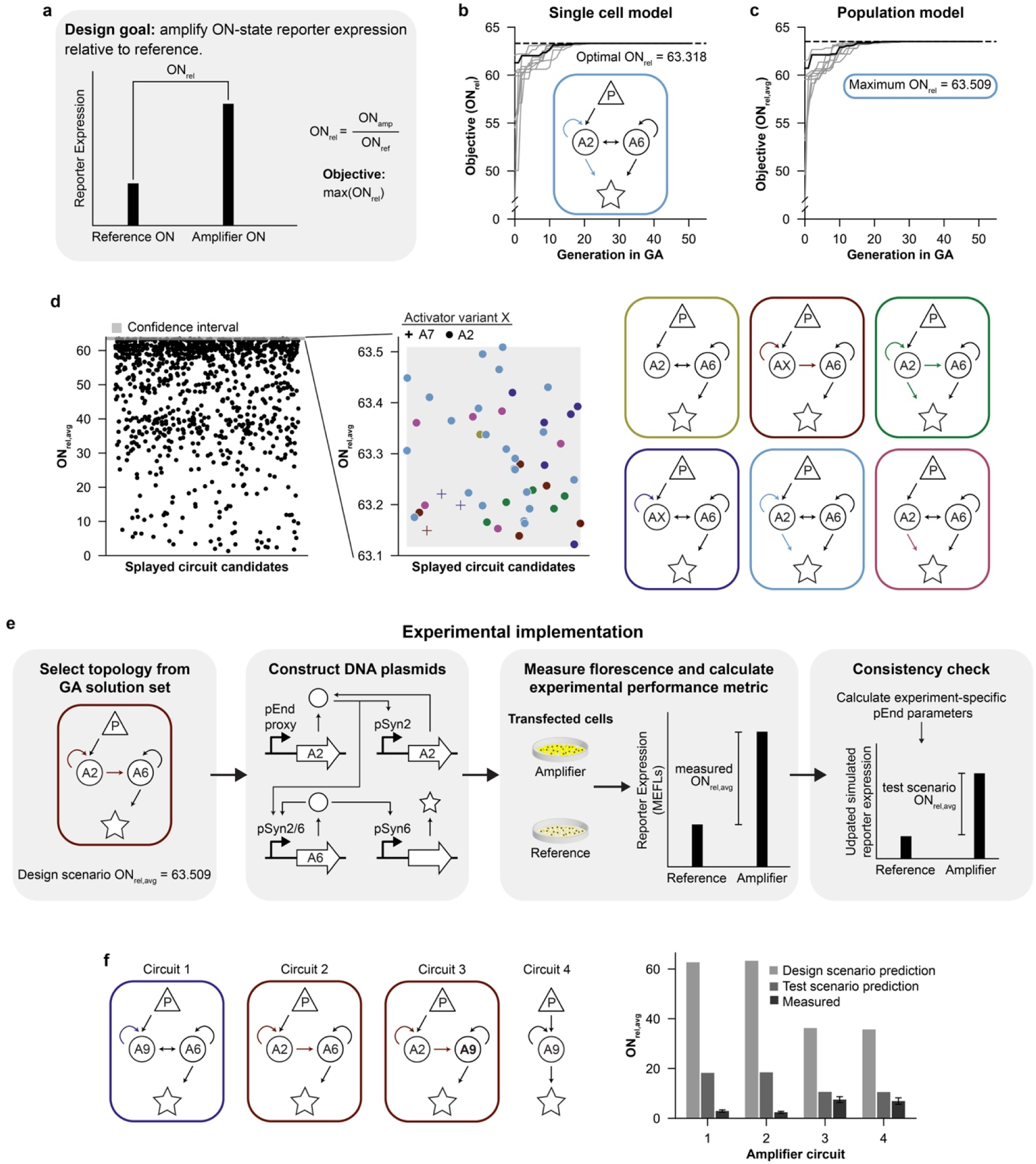
Optimal amplifiers are recovered with GCAD and evaluated in an experiment. **(a)** Schematic illustrating the design goal for the Amplifier test case translated into a quantitative metric for optimization (ON_rel_). (**b**) Optimization traces showing progression of ON_rel_ through each generation of the GA (single cell model for 10 different initialization seeds). The optimal ON_rel_ (dashed line) obtained from a combinatorial search and corresponding circuit. (**c**) Optimization traces showing progression of the population-level ON_rel_ (ON_rel,avg_) through each generation of the GA (20-cell population model for 10 different initialization seeds). The circuit with the maximum ON_rel,avg_ in each of the 10 seeds corresponds to the circuit in (**b**). (**d**) Full search space for the 20-cell population model. The ON_rel,avg_ values in the zoomed plot corresponding to the defined confidence interval are color-coded by the circuit designs on the right. (**e**) Workflow for experimental evaluation of circuits. A circuit is selected using GCAD based upon predicted ON_rel,avg_ (design scenario prediction), the circuit is constructed as DNA plasmids, which are transfected into cells, and the fluorescent reporter protein produced is quantified for each cell via flow cytometry. The reference case is also included in the experiment, which enables experiment-specific performance metric prediction (test scenario ON_rel,avg_), calculated post hoc using the reference circuit to evaluate the true ON and OFF states of the dox-inducible promoter which drives circuit activation. (**f**) Measured performance of selected circuits through transient transfection of HEK293FT cells, compared with design and test scenario predictions. Error bars for measured values indicate standard error of the mean, and each circuit was evaluated in biological triplicate. Data shown include transfected cells, as determined by a transfection control plasmid encoding a fluorescent protein. AX: synTFX-A (where X is the index of the synTF); P: endogenous promoter; R: reporter.

A key decision in simulation design is choosing how to represent a population of cells, and that choice is guided by the intended use of the simulation. With a focus on understanding our design algorithm, we opted to initially utilize a simplified, single cell genetic parts model (i.e., to simulate a single cell) to evaluate the performance of GCAD compared to a combinatorial search for the Amplifier test case. The single cell model does not account for population-level heterogeneity, for example due to differences in DNA uptake between cells (this is a natural feature of plasmid DNA transfection). However, this choice is useful for initial method evaluation because it is much less computationally expensive than a population model with heterogenous plasmid uptake, which we reserved for later consideration (**Methods**). In this tractable case, a combinatorial search of the design space is computationally feasible to determine the circuit with the optimal ON_rel_ value for comparison to that identified by GCAD.

We verified that our method finds the optimal ON_rel_ value and corresponding circuit. Here, the optimal circuit utilized the activators synTF2-A and synTF6-A, each at the highest possible component “dose” (75 ng of each plasmid), and yielded an ON_rel_ value of 63.318 (**Figure 2b**). The results were robust across 10 different initialization seeds with a set of hyperparameters obtained from preliminary manual tuning (Initial value, **Table 1**). The maximum ON_rel_ from each generation converged to the optimal value and corresponding circuit found by the combinatorial search for each of the 10 seeds (**Figure 2b**). Examining the range of candidate ON_rel_, we found that it matched the range for the combinatorial search space, implying that GCAD is adequately exploring the design space (**Supplementary Figure 1**). Furthermore, we found that the GA procedure only required the simulation of less than 1% of the combinatorial search space (total size: 1,679,760 topologies), thus highlighting the computational benefit of GCAD (**Supplementary Figure 2**). The GA circuit selection method with the single cell model successfully, consistently, and efficiently recovered the optimal solution for the Amplifier test case.

**Table 1.** GA hyperparameters, descriptions, and values for each test case, pre- and post-optimization.

| Hyperparameter | Description | Initial value: all test cases | Optimized value: Signal Conditioner<sup>1</sup> | Optimized value: Pulse Generator<sup>1</sup> |
| --- | --- | --- | --- | --- |
| GA population size | Size of the randomly generated population used to initialize GA | 52 | 200 | 200 |
| Initial ratio of 1- to 2-part circuits | Ratio defining the amount of 1- vs. 2-part circuits included in the population used to initialize GA | 0.5 | 0.25 | 0.25 |
| Crossover probability | Probability of crossover occurring in each generation of GA | 0.55 | 0.06 | 0.52 |
| Mutation probability | Probability of a mutation occurring in each generation of GA | 1 | 0.99 | 0.96 |

<sup>1</sup>GA population size and the population 1 to 2-part circuit ratio were fixed in the hyperparameter optimization.

We next expanded our model to simulate a population of cells for each circuit, to account for variation in DNA delivery (and the resulting activity of genetic circuits) between cells, and we re-examined circuit selection. The original modeling work^35^ used a 200-cell model, but this was computationally prohibitive during a GA search, where the full population is simulated each time a circuit topology is evaluated. By comparing simulations of circuits using a 20- or 200-cell model, we observed close alignment between objectives, indicating that a 20-cell representation was sufficient to capture the main features guiding circuit selection (**Supplementary Figure 3** and **Methods**). To perform a population-level GA search, the objective function was redefined as the average (i.e., arithmetic mean, which was selected to capture outlier behavior) ON_rel_ across the population, ON_rel,avg_. We ran the population-level GCAD with 10 different seeds, the same 20 cells in the population (i.e., a fixed distribution of DNA delivery variation), and the same initial hyperparameters (**Table 1**). For all seeds, the maximum ON_rel,avg_ converged to a similar value (**Figure 2c**) and the corresponding circuit was the same as the optimal circuit from the single cell model search (**Figure 2b**). Notably, the single cell model search recovers one optimal solution and circuit, while there is not a unique, optimal solution for the population model search.

We next examined the source of variability for the same circuit between seeds. Within each cell in the population, the copy number of parts can vary because our model specifies the dose of different plasmids based on the order in which the corresponding parts are added to the network. Thus, a single circuit with parts A2 (synTF2-A) and A6 (synTF6-A) can have two possible combinations of relative dosages (e.g., A2^high^A6^low^ or A6^high^A2^low^) for each run (i.e., seed) of a simulation. Thus, although we are not testing different populations of cells in each seed, there is still variability in expression of the circuits from seed to seed, leading to variability in ON_rel,avg_. We expect that the variation in ON_rel,avg_ values would be larger if we captured the heterogeneity in *in vitro* cell populations across experimental replicates by drawing cells to create multiple distinct populations. For either source of variation, it is relevant to consider an ensemble or *set* of solutions at or near the maximum ON_rel,avg_. To determine if the GCAD search is robustly recovering the same ensemble of circuits, we must define a confidence interval (CI) on the objective value that captures the spread or variation due to population variation and determine if the same set of circuits appear within this confidence interval across multiple GCAD runs. To this end, we investigated whether differences in circuit design produce detectable differences in ON_rel,avg_ when accounting for population variation, how such differences affect the GCAD search, and whether this search is robust.

We defined a confidence interval (CI) for the most optimal population-level average metrics found for each test case to specify a set of solutions that would be indistinguishable from optimal when accounting for population variation between experimental replicates. Any circuit within the CI could therefore be considered an equally optimal design (e.g., to be selected for experimental implementation). To define a CI, we first selected circuits with ON_rel,avg_ values in a distinct cluster close to the maximum value—62.600 ≤ ON_rel,avg_ ≤ 63.509. Then, we simulated the selected circuits with 10 different manifestations of the 20-cell population model and calculated the standard error in ON_rel_ across the simulations for each circuit. We defined the CI as follows:

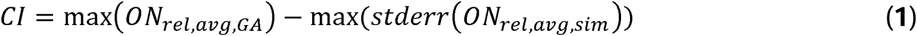

Here, max(ON_rel,avg,GA_) is the maximum ON_rel,avg_ from the GCAD search with a selected seed, and max(stderr(ON_rel,avg,sim_)) is the maximum standard error in ON_rel,avg_ across the simulations for the selected circuits for all 10 manifestations of the population model. Importantly the CI can be calculated after the GCAD search and does not require multiple GA runs. We considered any ON_rel,avg_ values within this CI as indistinguishable from optimal due to population variation. A separate question that the user must evaluate is whether the variation within this ensemble is either biologically meaningful or relevant to the design goal. We expect that for different population instances, a robust GCAD search will return the same ensemble of circuit designs within the confidence interval.

To validate the robustness of the 20-cell population-model based search with CI, we must evaluate whether the CI calculated using the 20-cell model was sufficient to capture the variation compared to that of the 200-cell model. We used the same strategy as for the 20-cell model to calculate a CI for the 200-cell model. The 200-cell CI was included within the 20-cell CI (**Supplementary Figure 4a**), with a smaller range than that of the 20-cell model. This result suggests that the 20-cell model is sufficient to predict ON_rel,avg,_ particularly for high-performing topologies. We additionally simulated topologies from the 20-cell model GCAD search space that had high ON_rel,avg_, mid-range ON_rel,avg_, and low ON_rel,avg_ with the 200-cell model to compare alignment between ON_rel,avg_ for the two models. We found good alignment between ON_rel,avg_ for the two population sizes (**Supplementary Figure 4b**), indicating that the 20-cell model can sufficiently predict ON_rel,avg_ for topologies with a range of performances. Overall, this gave us confidence that if GCAD was run with the 200-cell model, it would likely explore similar candidate circuits to those explored with the 20-cell model to converge on a similar solution set.

We found that all the circuits within the CI were one of 6 topologies with positive and auto-regulation (**Figure 2d**), containing maximal or near maximal doses of the activator synTF6-A and either synTF2-A or synTF7-A activators. SynTF6-A, synTF2-A, and synTF7-A are the 3 strongest transcriptional activators in the set of synTF-As used here, so this finding aligns with our expectations that these components would produce the largest amplification in reporter expression. Different manifestations of all the circuits within the CI, i.e., circuits having the same topologies but comprising different activator variants or component doses, also appeared outside of the CI; they generally had at least a mid-range predicted ON_rel,avg_ value (≥ 30). These different manifestations of CI circuits either contained the same synTF-A variants as one of the optimal circuits (with lower doses of the synTF-As) or one of the other, weaker, synTF-A variants (generally in combination with either synTF6-A, synTF2-A, or synTF7-A). Within the CI, the topology indicated in blue (**Figure 2d**) was by far the most common, followed by the topology indicated in pink, indicating that autoregulation of both synTF-As and activation by the other synTF-A enables the highest predicted amplification of gene expression. Outside of the CI, the topology indicated in green was by far the most common, followed by the topology indicated in blue (**Figure 2d**), suggesting that autoregulation and dual activation of the reporter can enable at least mid-range gene expression amplification even with weaker synTFA-s or lower component doses. Overall, designing an amplifier with ON_rel,avg_ at or near the maximum attainable with the synTF-As used here required not only two of the strongest activator variants at or near the maximum possible dose, but also either dual autoregulation, bidirectional activation of each amplifier, or dual activation of the reporter (and generally a combination of these regulatory motifs). Circuits that did not meet all three of these criteria exhibited some level of amplification but did not operate at or near the maximum ON_rel,avg_.

Finally, we selected and experimentally evaluated a set of amplifier circuits identified using GCAD to evaluate their behavior in cell culture. Thus far, our comparison between the combinatorial search and the 200-cell model provided confidence that GCAD robustly finds the optimal ensemble of circuits as modeled. However, it was still possible that the models will not predict the empirical circuit performance; as noted, the original model was designed to provide mechanistic insights,^35^ and GCAD considers some conditions (e.g., plasmid doses) where the model’s predictive capabilities were not explicitly validated during model development. Towards the goal of exploring how such considerations guide use of GCAD, we selected circuits as candidates for experimental evaluation from two clusters of ON_rel,avg_ values from the Amplifier search with the population model—one near the maximum value, representing a best performing circuit from the CI-ensemble, and another around two thirds of the maximum representing sub-optimal circuits.

We constructed a set of DNA plasmids encoding each circuit topology (**Supplementary Figure 5**), as well as a reference case representing the “unprocessed” output of an endogenous promoter which motivates our design goals. We used a doxycycline (dox)-inducible Tet-On system^37^ to serve as a proxy for the behavior or an endogenous promoter (pEnd), enabling us to control the activation states used to probe circuit performance. We performed a dox dose-response characterization using this system to drive expression of a reporter protein, from which we selected dox concentrations to represent the OFF and ON states of pEnd (**Supplementary Figure 6**). These values were selected to represent a promoter with some non-zero basal leak (which is a commonly encountered behavior) rather than a more trivial zero-leak OFF state (for which fold-induction cannot be calculated). For subsequent circuit evaluation experiments, the regulators of a circuit that are driven by the endogenous promoter (e.g., a synTF-A) were placed under control of the same Tet-On system, and the circuit was evaluated by inducing with the dox concentrations corresponding to OFF and ON values of pEnd. In those experiments, the plasmids mediating dox-inducible expression of the reporter served as the reference circuit (activated using the concentration of dox corresponding to the OFF and ON states of pEnd).

Circuits were functionally evaluated by transfecting the specified doses of plasmids encoding each component into HE293FT cells and inducing circuits with dox as described above. After allowing time for reporter expression (∼2 days, **Methods**), fluorescence of individual cells was measured by via flow cytometry, and the measured ON_rel,avg_ values were calculated for each amplifier (**Figure 2e**). Each amplifier circuit exhibited a measured ON_rel,avg_ greater than 1, indicating amplified reporter expression (**Figure 2f**); however, the predicted ON_rel,avg_ values did not align well with the experimentally measured values, prompting further investigation.

We next explored possible reasons as to why the model overestimated the degree of amplification our test circuits would exhibit. First, we performed a consistency check; we used the reference circuit to check whether the scenario used to guide model predictions (design scenario) and the scenario implemented (test scenario) match. The observed reference circuit behavior differed from that used to guide circuit design; in this case, the reference circuit exhibited a higher fold-induction than expected. Such variations between experiments are common (due to differences in cell division rate, transfection efficiency, and other factors that are hard to experimentally control); this illustrates the necessity of including such a reference circuit. Using the reference circuit outputs to calculate the OFF and ON states actually employed to induce the test circuits (using an empirical calibration curve; **Supplementary Figure 6**), we re-simulated circuit behavior. The updated predictions (test scenario) (**Figure 2f**) were much closer to the measured values for Amplifiers 3 and 4, but they still overestimated ON_rel,avg_ values generated by Amplifiers 1 and 2.

Thus, we continued our analysis to identify the source of discrepancy. Another key clue is that Amplifiers 3 and 4 exhibited higher measured ON_rel,avg_ values than Amplifiers 1 and 2, contrasting with the predicted trend. Upon inspection, we noted that Amplifiers 1 and 2 both utilize synTF6-A, which is the most potent of the synTF-A variants in this set. Prior work has suggested that some synTF-A variants can exhibit transcriptional squelching, whereby excess transcription factors create a sink that sequesters transcriptional co-factors away from the DNA-bound transcription factors.^35^ Squelching is not currently represented in the ODE model; prior usages of these parts simply avoid this squelching regime by restricting synTF-A levels^6^, but no such constraints were placed upon GCAD during the design phase. Altogether, these observations suggest a likely explanation as to why GCAD-selected circuits perform in unexpected ways when high levels of a potent factor (synTF6-A) are deployed. Furthermore, this mechanistic insight provides testable hypotheses as to how this disconnect could be avoided through model refinement (e.g., by experiment-guided refinement of the ODE model to capture squelching) and/or through evolving the constraints provided to GCAD during the design phase (e.g., avoiding high levels of synTF-6A or omitting this part from consideration altogether). For the purposes of this study, our comparison of predictions to experiments is really a test of the predictive capability of the genetic parts model, not the GCAD framework. Overall, this test case illustrated the feasibility of using GCAD for circuit design, while providing important insights into how understanding ODE model accuracy across operating regimes may guide a choice of the most useful GCAD constraints.

### Test case 2: multi-objective amplification of output gene expression with increased inducibility

We describe our second test case as a Signal Conditioner, for which the design goal is to both amplify the ON-state and increase the fold induction (the ratio of output gene expression in the ON-state to that in the OFF-state). For the quantitative design specification, we used ON_rel_, as in the Amplifier test case, and we defined the second objective to be the ratio of the Signal Conditioner fold induction (FI) over the reference case FI, FI_rel_. Circuits having ON_rel_ and FI_rel_ values that are each greater than 1 would satisfy the design goal, and we maximized both in a multi-objective optimization within the GA (**Figure 3a**). The design space was the same as that used in the Amplifier test case—all transcriptional activators and repressors with all possible doses (within the original range) and interactions were considered.

**Figure 3.**
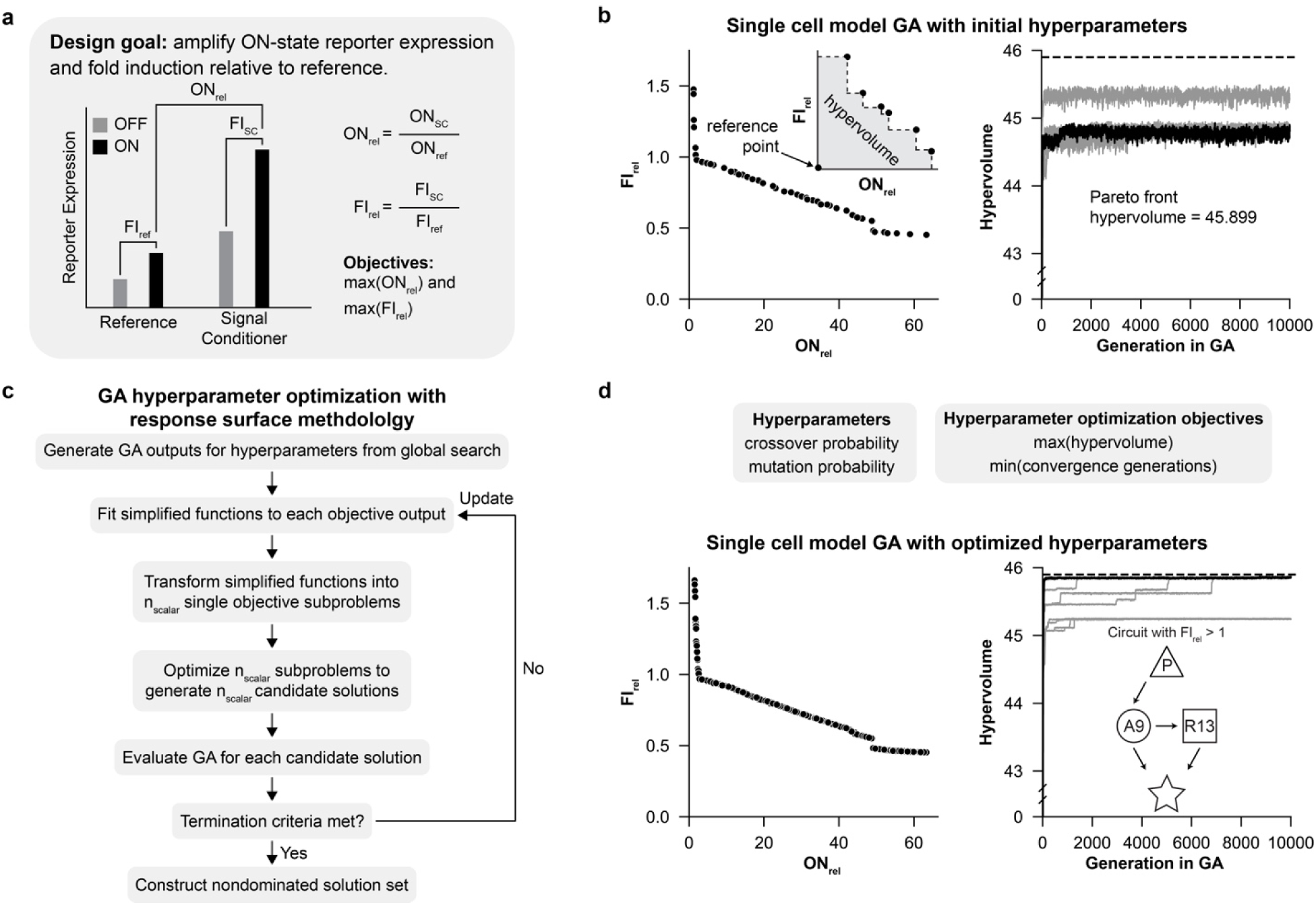
GCAD successfully recovers the majority of Pareto optimal solutions for the Signal Conditioner. **(a)** Schematic illustrating the design goal for the Signal Conditioner test case posed as a set of quantitative objectives for optimization: relative amplification, ON_rel_, and relative fold induction, FI_rel_. (**b**) GCAD identification of an initial Pareto front of the tradeoffs between objective functions, using heuristically determined hyperparameters (left). The hypervolume quantifies the coverage of the non-dominated solution set in multi-objective optimization (left, inset plot). The hypervolume converges near the optimal hypervolume (dashed line) determined by the combinatorial search for 10 different initialization seeds (grey) (right). (**c**) Hypervolume optimization method using response surface methodology (RSM). RSM fits simplified functions to each of the hyperparameter optimization objectives to decrease the cost associated with running the full simulations. (**d**) GCAD identification of a Pareto front of the tradeoffs between objectives using the RSM optimized hyperparameters (left). The hypervolume converges to near the optimal found by the combinatorial search (dashed line) for 10 different seeds (grey) (right). GCAD solutions with FI_rel_ > 1 had the same circuit structure with different component doses.

We first assessed the performance of the single-cell model GCAD for the Signal Conditioner compared to a combinatorial search. Leveraging multi-objective optimization, we found the set of Pareto optimal solutions for which neither objective can be improved without worsening the other. Hypervolume^38^ quantifies the area enclosed by the Pareto front and a specified reference point, in this case (0, 0), allowing us to compare solution sets between GCAD runs (**Figure 3b**). As in the previous test case, we ran GCAD for 10 different initialization seeds using hyperparameters determined by preliminary manual tuning (**Table 1**). For nine out of ten seeds, the hypervolume approached values between 44.7 and 44.9 by generation 600, and it fluctuated between these values for the remainder of the 10,000 generations (**Supplementary Figure 7**). The fluctuations in hypervolume are due to the constraint of GA population size, which prevented some solutions along the Pareto front from advancing to the next iteration, and the existence of local optima with low crowding scores, which can take the place of true Pareto solutions as the non-dominated sorting algorithm prioritizes candidates in lesser crowded region to promote diversity. The hypervolume fluctuated as the GA found first one set of optimal solutions and then another since the small solution set cannot fully cover the range of Pareto optimal solutions. The seed with the maximum hypervolume approached values between 45.2 and 45.4 by generation 400 and similarly fluctuated between these values for the remainder of the 10,000 generations. The maximum hypervolume obtained by GCAD was 1.1% less than that of the Pareto front obtained from the combinatorial search (45.899), since not all Pareto optimal solutions could be found simultaneously (**Figure 3b**). Given the fluctuations in hypervolume for the set of hyperparameters from the Amplifier test case, we chose to investigate whether hyperparameter tuning for this test case could enable consistent convergence at the Pareto front hypervolume.

We hypothesized that better calibration of the GA hyperparameters could enable faster convergence, decreasing computational expense, so we performed a hyperparameter optimization to test this hypothesis. We utilized response surface methodology^39, 40^ (RSM) to find hyperparameters that simultaneously maximized the hypervolume and minimized the number of generations taken for GCAD to converge on the final hypervolume. RSM is a multi-objective optimization approach in which a less computationally expensive, surrogate function is fit to each of the objectives to avoid the need to run the more costly simulation—in this case, the GA procedure in GCAD—for each set of candidate hyperparameters (**Figure 3c**, **Methods**). We used a Gaussian radial basis function as a surrogate function for hyperparameter optimization, although we note that polynomial regressions are also commonly used.^39^ To enable simultaneous recovery of more of the Pareto front, we increased the GA population size from 52 to 200. We optimized the probability of crossover and mutation and fixed the initial ratio of 1 to 2-part circuits in the population at 0.25.

We next ran GCAD for 10 seeds with the most optimal crossover and mutation probabilities found in the hyperparameter optimization (**Table 1**), which resulted in consistent hypervolume convergence at different rates. The six seeds with the maximum hypervolume reached hypervolumes within 0.1% of the Pareto front hypervolume by generation 90 for three out of six seeds, by generation 1400 for one of seeds, and by generations 5100 and 7000 for the other two seeds, respectively. For the other four out of ten seeds, the hypervolume converged to values within 1.5% of the Pareto front hypervolume by generation 300 and 600 for two of the four seeds, respectively, and by generation 1300 for the other two seeds. In all cases, the final solution set has better coverage of the Pareto front, and several additional solutions with FI_rel_ greater than 1 were found, some of which had a FI_rel_ greater than the maximum value obtained with the initial hyperparameters (**Figure 3d**). These improvements were due to both the increase in GA population size and the optimization of crossover and mutation rates. If only the population size is increased, only 2/10 seeds converged to within 0.1% of the Pareto front hypervolume, indicating that the hyperparameter optimization significantly increased convergence (**Supplementary Figure 8**). Solutions with FI_rel_ greater than 1 were all the same circuit differing only in component doses, indicating that the recovered topology has the potential to amplify ON_rel_ and FI_rel_, and it is possible that this topology could even better achieve the design goal if constructed with a different set of parts (i.e., new parts not included in our design space). Comparison of the GA Pareto front with that generated in the combinatorial search indicated that there were 4 optimal solutions with FI_rel_ greater than 1 that were not recovered with the GA; these had FI_rel_ values of 1.03, 1.33, 1.35, and 1.38. These “missed” solutions were each close to another Pareto optimal solution that was recovered by GCAD, within 0.0062 FI_rel_ and 0.085 ON_rel_ (overlapping on the plot in **Supplementary Figure 9a**). As in the Signal Conditioner test case, GCAD recovers solutions across the combinatorial design space, suggesting the algorithm balances exploration convergence to Pareto optimal solutions (**Supplementary Figure 9b**). In addition, we found that GCAD required significantly fewer topology simulations than did the combinatorial search (less than 0.5% of all possible topologies; **Supplementary Figure 2**) while attaining high hypervolumes compared to that of the true Pareto front (roughly 97%). This reduction in computational efforts without compromising solution quality reinforces the potential of extending GCAD to more complex dynamics.

We performed circuit selection with the population model to select candidates for experimental evaluation using an analogous workflow to that of the Amplifier test case. In this case, the circuits that best satisfied the design goal had FI_rel,avg_ and ON_rel,avg_ values near 1 (**Supplementary Note 3**, **Supplementary** Figures 10**, 11**). Given this modest projected performance and expected experimental variability, we concluded that empirical evaluation would be unlikely to differentiate Signal Conditioners’ performance from that of the reference circuit. Therefore, we chose not to pursue further analysis of the Signal Conditioner. This case illustrates how GCAD can provide confidence that, sometimes, an idea is not worth pursuing with the parts available, enabling the designer to repurpose time and resources.

### Test case 3: generation of a pulse in output gene expression

The final design goal we evaluated is to produce a pulse in output (reporter) expression, which we refer to as the Pulse Generator. This is the first test case to focus on producing desired dynamic behavior. We used two different metrics in the design specification to quantify whether the reporter expression exhibited a pulse relative to the reference case (reporter_rel_) (**Figure 4a**). First, we quantified the magnitude of the pulse as the peak prominence of the reporter_rel_ (prom_rel_). Prominence is defined as the difference in maximum value of the reporter_rel_ and value of the reporter_rel_ at the final simulation time point. Prom_rel_ was set equal to 0 if the reporter_rel_ time series simulation did not contain a peak (i.e., the end value was the maximum value). A second metric was the time at which the pulse occurs (t_pulse_), defined as the time at which the reporter_rel_ reaches the peak, or the maximum simulation time if reporter_rel_ does not contain a peak. The objectives for this test case were to maximize prom_rel_ while minimizing t_pulse_. We again chose a design space including the full set of synTFs and promoters to test GCAD’s capabilities on a complex search space.

**Figure 4.**
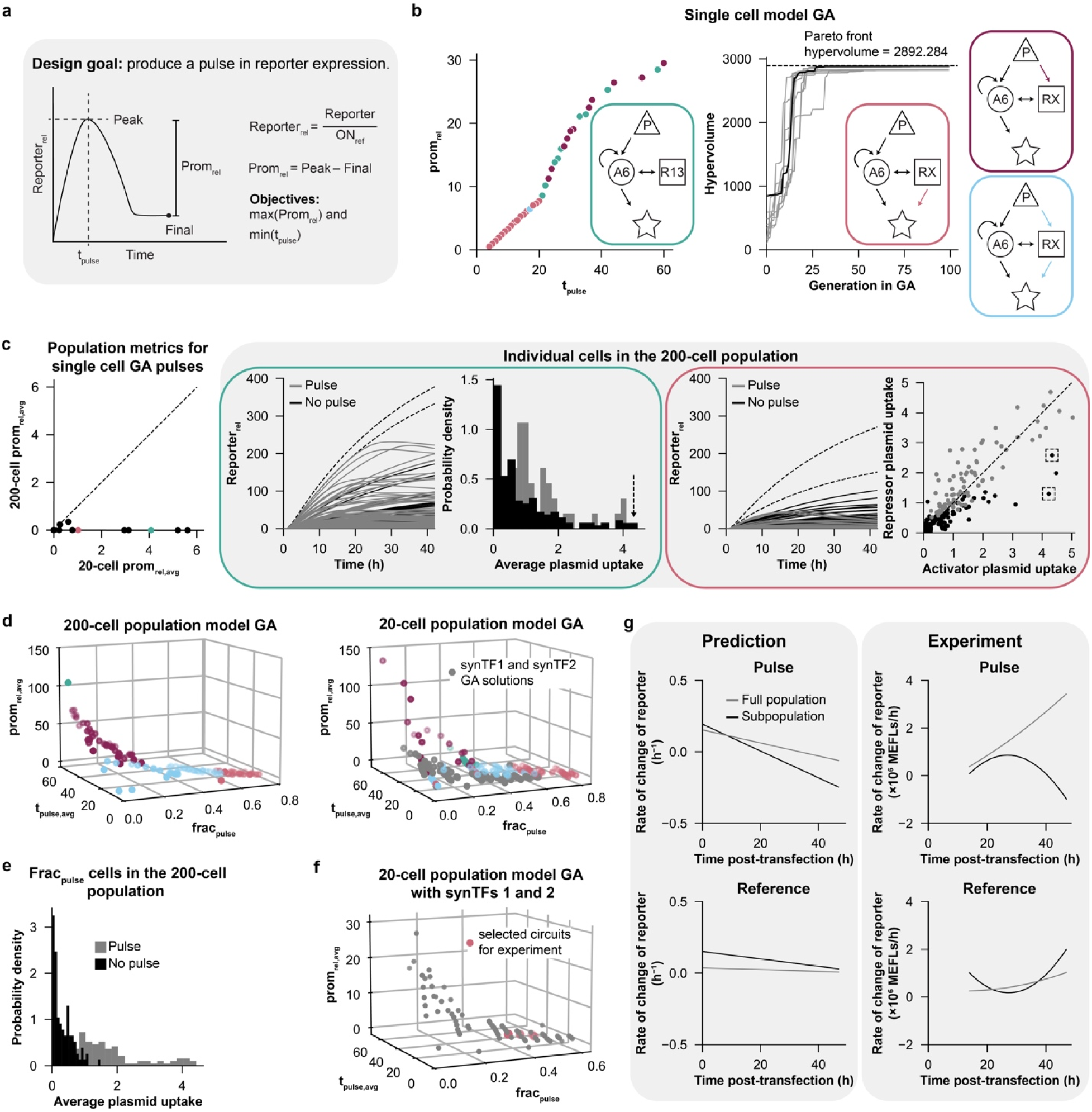
The Pulse Generator test case underscores the importance of the choice in optimization objectives. (**a**) Schematic showing the design goal for the Pulse Generator test case translated into a set of quantitative metrics for optimization: the time to pulse, t_pulse_, and relative prominence, prom_rel_. (**b**) The single cell model GCAD solution set for a representative seed (left) and the hypervolume progressions across 10 different seeds compared to the optimal hypervolume from a combinatorial search, finding three distinct circuit architectures (right, circuits color coded). (**c**) Performance evaluation of found circuits. While the population-level average for the 20-cell model produces pulse behavior, only a few cells in the 200-cell model produce a high reporter_rel_ and they do not produce significant pulsing behavior (teal/blue color references circuit diagrams in **b**). Each cell is classified as pulsing (grey) or not pulsing (black) based on quantitative detection of a peak, which may not be visually apparent; in both insets, the dashed traces exemplify cells with high reporter expression but no pulse. Within the teal circuit inset, the bar of the histogram identified with the dashed arrow (right, highest average plasmid uptake) corresponds to the traces with dashed lines (left). Within the blue circuit inset, cells identified in dashed squares (right) correspond to traces with dashed lines (left). (**d**) Comparison of Pareto fronts for 200-cell and 20-cell model GCAD runs incorporating a new metric, frac_pulse_, the fraction of individual cells in the population that produce a pulse. This new search recovered an additional circuit (light blue). Solutions using only synTF1 and synTF2 are highlighted (grey). (**e**) Pulse behavior as a function of plasmid uptake. Cells with a higher average plasmid uptake pulse (top 30%), while cells with low average plasmid uptake (lower 70%) do not pulse; shown is a representative example of the circuits in **d** for the 200-cell model. (**f**) 20-cell model GCAD runs incorporating frac_pulse_ with a subset of the parts library—including synTF variants 1 and 2 only. The topologies selected for experimental evaluation are indicated. (**g**) Comparison of predicted and experimentally measured Pulse circuit performance for a selected circuit (Pulse 1 circuit). To summarize the behavior of ensembles of single cell traces (200-cell model) and focus on signatures of pulse behavior, we fit a polynomial to the reporter levels for all cells and calculated the instantaneous rate of change (first time derivative) from the polynomial fit (left). This analysis was performed for both the full population of cells and for a subset predicted to exhibit a pulse. Pulse circuit 1 was also experimentally evaluated following the same methodology as the Amplifier test case, and reporter levels were processed, both for the full population and for a subset identified by high plasmid uptake (top 30%, based on a transfection control plasmid; right).

First, we assessed the performance of GCAD with the single cell model for the Pulse Generator, in comparison to a combinatorial search. The combinatorial search provided a set of Pareto optimal solutions, from which we calculated the hypervolume for comparison to the GCAD solution set. We calibrated the GA hyperparameters using RSM optimization (**Figure 3c, Methods**) and ran GCAD for 10 different initialization seeds with the most optimal set of hyperparameters found (**Table 1**). The solution set consisted of pulses with t_pulse_ between 4 and 60 hours and prom_rel_ between 0.5 and 30. For four out of ten seeds, the hypervolume converged to values within 2.5% of the Pareto front hypervolume by generation 80. The six seeds with the maximum hypervolume converged to hypervolumes within 0.4% of the Pareto front hypervolume by generation 80 (**Figure 4b**). There were four different topologies among the Pareto optimal solutions (**Figure 4b**); interestingly, the most common topology (dark blue) was similar in structure to an incoherent feedforward loop—a naturally occurring network motif that is known to produce a pulse in gene expression.^41, 42^ In addition to this known motif, the found topology also had activator autoregulation and repressor regulation of the activator, an unintuitive addition that highlights the potential utility of a semi-automated design approach. The other topologies (teal, light blue, and maroon) had very similar structure but variability in the regulation between the promoter and the repressor and the repressor and the reporter. Comparison of the GA Pareto front with that generated in the combinatorial search indicated that there were 5 optimal solutions that were not recovered with the GA (**Supplementary Figure 12a**). The explored GCAD search space contained solutions across the combinatorial design space, demonstrating that it explored diverse regions of the design space to converge on most of the Pareto optimal solutions for this multi-objective optimization problem (**Supplementary Figure 12b**).

We next examined how Pulse Generator circuit performance depends on the size of the population model. When comparing simulations of 200-cell and 20-cell models of circuits in the GCAD solution set, there were discrepancies in the average population behavior, including both t_pulse,avg_ and prom_rel,avg_ values. Notably, when only considering the population-level average behavior, there were several cases in which the 200-cell model prom_rel,avg_ was zero, while the 20-cell model prom_rel,avg_ was significantly greater than zero, indicating that reporter_rel,avg_ for the 20-cell model exhibits a pulse, while this same metric derived from the 200-cell model does not exhibit a pulse (**Figure 4c**). For each case where the 200-cell prom_rel,avg_ was equal to 0, examining the reporter_rel_ time series for individual cells from 200-cell model simulations revealed that several cells pulsed, even when the population on average exhibited no substantial pulse (**Figure 4c**). Upon inspection, we noted that some non-pulsing cells in the 200-cell model had a much higher maximum reporter_rel_ than did the pulsing cells in each case, which skewed the population-scale reporter_rel,avg_ and masked the pulsing of some cells (**Figure 4c**, dashed curves).

To better understand differences between pulsing and non-pulsing cells, we next compared the distributions of average plasmid uptake for each sub-population. We first examined cells exhibiting high reporter output but no pulse (**Figure 4c**, teal and blue boxes, dashed traces). In the case of the teal circuit, the cells with the highest maximum reporter_rel_ took up the highest average amount of plasmid (**Figure 4c**, teal box). For the most frequently found circuit (blue), the cells with the highest maximum reporter_rel_ again took up a high average amount of plasmid, but they also took up more activator plasmid than repressor plasmid (**Figure 4c**, blue box, dashed boxes). The cells that produced a pulse could not be distinguished from those that did not pulse based upon average plasmid uptake alone. Compared to the 20-cell model, the 200-cell model better sampled cases in which high plasmid uptake led to high reporter output and no pulse (and these outliers suppressed the performance metric, prom_rel, avg_, for the 200-cell model; **Figure 4c**, left panel). However, since there nonetheless existed individual cells exhibiting pulses in both models, this analysis demonstrates how choice of objective for optimization can influence solution recovery. In this case, the t_pulse,avg_ and prom_rel,avg_ objectives are perhaps useful for optimizing for pulse behavior that is robust across a population of cells, but these objectives may be less appropriate for identifying circuits that produce a pulse within at least a subset of the population. This analysis also highlights how population-level objectives (especially arithmetic mean metrics) may be biased (or confounded) by the behavior of rare cells.

We next investigated whether an objective less affected by outliers (e.g., cells within tails of the plasmid uptake distribution) could be used to identify pulse behavior in a subset of a population. To that end, we defined a new objective—the fraction of cells that produce a pulse (frac_pulse_). We used this objective in addition to t_pulse,avg_* and prom_rel,avg_*, quantified as the average t_pulse_ and prom_rel_ across only the cells in the population that produce a pulse. We then generated GCAD search and solution sets using the 20-cell model and the 200-cell model in this 3-dimensional multi-objective space (**Figure 4d**). The solution sets were well-aligned between the two population models; the ranges of performance metrics for the most optimal solutions found was similar, and resulting topologies were the same. Interestingly, within these circuits, average plasmid uptake could be used to distinguish cells that produced a pulse from those that did not (**Figure 4e**), as pulsing cells had a higher average plasmid uptake than non-pulsing cells. This outcome has practical utility, as it is possible to physically isolate cells with higher plasmid uptake, for example by using fluorescence activated cell sorting to enrich for highly transfected cells (e.g., using a transfection control plasmid expressing a fluorescent protein marker). Overall, careful choice of objective function and its relationship with population heterogeneity can improve the search for useful circuit designs, and GCAD offers a flexible framework for exploring these effects.

Finally, we experimentally evaluated GCAD-designed Pulse Generator circuits. To avoid challenges observed in the first test-case when the design space exceeded the experimental characterization space, we restricted this GCAD design step to use only synTF1 and synTF2 activators and repressors, since due to a quirk in their DNA recognition sequence, they are particularly effective repressors, and the interaction of repressors and activators based upon these synTFs was explicitly experimentally observed and modeled in the original study^35^. Comparing 20-cell GCAD runs using all synTFs or using only these two, we found that the solution sets spanned a similar range of objective values, but circuits using only synTF1 & synTF2 slightly underperformed the broader search in each metric (**Figure 4d**). We prioritized circuits from the synTF1 and synTF2 solution set with t_pulse,avg_ between 10 and 24 h to maximize the probability of observing such dynamics using our experimental setup. Within this set of solutions, we further downselected to four circuits having both a high fraction of pulsing cells, frac_pulse_, and high prominence, prom_rel,avg_ (**Figure 4f**).

To set expectations for the experiment, we analyzed the simulations of individual cell pulse behavior for the selected circuits as a function of plasmid uptake (200-cell model). At around 60% average plasmid uptake, all cells in the simulated population produce a pulse for each of the selected pulse circuits (**Supplementary Figure 13**). Therefore, filtering the cell population based on plasmid uptake could select for a subpopulation of cells that exhibit pulse behavior. We conservatively selected a plasmid uptake percentile of 70% as a threshold for filtering the “high DNA uptake” cell population, and we compared the time series of the individual cells in the full 200 simulated cells population to those of the filtered subpopulation. Because the long half-life of the reporter protein could preclude a substantial decrease in reporter fluorescence during the experimental time frame, we instead quantified the instantaneous rate of change in reporter in order to detect a pulse in expression rather than fluorescence. To calculate the rate of change, we fit an interpolated polynomial to the reporter_rel_ at every time point for each cell and calculated the derivative. A positive derivative indicates the expression of the reporter is increasing, and a negative derivative indicates the expression of reporter is decreasing. So, if the derivative transits from a positive value, through zero, to a negative value, this indicates that the production of reporter has peaked, and the expression exhibits a pulse. For the reference circuit we did not expect a pulse, and indeed both the simulated subpopulation and full population exhibited rates of reporter change that were positive and approached but did not cross zero, indicating a plateau rather than a peak (**Figure 4g**). For the circuits designed to produce a pulse, the rates of reporter change were distinct between the simulated full population and the subpopulation. The derivative of the reporter crossed zero (i.e., reporter peaked) around at around 20 h for the subpopulation, while the full population did not exhibit a reporter peak until around 33 h and since the derivative of the reporter never passed zero by much in this case, the full population peak was poorly defined (representative example, **Figure 4g**, other circuits, **Supplementary Figure 14**). Overall, this analysis suggests that evaluating the rate of change of reporter levels over time is useful for identifying and even characterizing pulse behavior in this system.

We next experimentally tested the pulse and reference circuits. We used a similar experimental implementation as in the Amplifier test case (**Figure 2e**), but in this case we measured single cell fluorescence at 7 time points after DNA transfection (14-46 h). We calculated the instantaneous rate of change of reporter level at each time point in the experimental data, as was done for simulation data. To identify the subset of cells with the highest amount of DNA uptake, which we expect to be more likely to exhibit a pulse, we used post-hoc analysis guided by expression of a transfection control plasmid (subset is top 30%, as used in simulation analysis). Compared to the simulations, the rate of change in expression for the reference circuit exhibited more variation over time but did not cross zero, indicating no peak in expression for either the full or subpopulation data (**Figure 4g**). Such variations within the positive rate of production regime can arise due to changing rates of cell division and metabolism over time. For the circuit designed to pulse, we successfully detected a peak in expression for the high DNA uptake subpopulation and not for the full population, qualitatively confirming our expectations. For the subpopulation, the rate of change in reporter expression initially increased and then decreased, crossing 0 at about 41 h and reaching a negative value by the end of the time series. In the full population, the change in reporter expression increased and remained positive, indicating no clear pulse (**Figure 4g**). Overall, GCAD was useful for iteratively and successfully designing a circuit which produced a measurable pulse in a subpopulation of cells. This is particularly impressive given that the parts in this design space appear to be (even from our simulation analysis) not necessarily well-suited to generating a pronounced version of this dynamic behavior. This success also supports our adjustment of the design criteria used in the GA search to incorporate population effects and explicitly select circuits that produce a larger subpopulation of cells exhibiting the desired behavior.

## Discussion

In this study, we developed and evaluated GCAD—a framework for semi-automated design of mammalian genetic programs. GCAD extends the horizon of model-guided design beyond the limits of a researcher’s ability to propose candidate programs to meet a performance goal based upon intuition, prior experience, or iterative experimentation. As part of this work, we proposed conceptual framings for posing design specifications (which may be qualitative and can include either steady-state or dynamic behaviors) as a quantitative objective or set of objectives that enable use of optimization to guide circuit selection. Finally, we employed wet lab experimental evaluation of GCAD-designed circuits to both guide refinement of GCAD and its usage and to validate its utility for designing genetic programs.

Our experimental evaluation of circuit solutions shed important light on the connections between ODE model development and the nature of resulting GCAD-selected designs. As noted at the outset of this investigation, we opted to use an existing ODE model describing a library of synTF parts and promoters that are well-suited to developing GCAD, even though (i) the original model was designed to provide mechanistic insights rather than prediction and (ii) some parts and interactions (e.g., competition between synTF-A and synTF-R pairs) were explicitly experimentally evaluated, while other interactions were not.^35^ One option would be to start by reducing these model uncertainties, for example by empirically characterizing more interactions and behaviors and by improving model confidence by employing rigorous parameter sensitivity analysis, model reduction, and/or uncertainty reduction-motivated experimentation.^5^ In particular, the original ODE model focused on explaining long-term behavior, such that the dynamic behaviors of the parts were not fully characterized; such model refinement could improve predictions of dynamic circuit behaviors. While employing high confidence models is preferred, there is also a substantial cost to such model improvements, and thus we opted to proceed with a model having some utility but known limitations in order to explore how this impacted GCAD-guided model selection.

A key lesson from the Amplifier case is that allowing GCAD to consider parts of design space, such as regimes of component dose, that have not been empirically evaluated can introduce new phenomena that lead to poor quantitative predictions. In this case, deployment of synTF6-A at very high levels induced apparent squelching in empirical tests, but since squelching was not encountered in prior work using lower synTF6-A doses, it was not modeled; this led to disconnect between predicted and observed behavior. One take away may be that establishing empirical limits on component usage (bounds on which behavior is well-captured by a model) may help to avoid this disconnect. This view is supported by the fact that GCAD-guided selection of Amplifier circuits employing components that do not exhibit squelching led to good qualitative and quantitative agreement between predicted and observed performance.

The Signal Conditioner test case exemplifies a valuable (if less glamorous) usage of computer-aided design—informing the user when not to proceed with experimental implementation. In this case, GCAD provided a high confidence conclusion that no “good” circuit design exists for meeting the design goal for this case. Simply avoiding wasted experimental effort is valuable. However, that conclusion is restricted to the existing design space. Thus, in practice, hitting such a wall could motivate one to explore whether new parts (e.g., hypothetical parts with parameters beyond or other than those in the empirical set, interactions that differ qualitatively or quantitatively from those which were measured, etc.) could improve circuit performance. Although a systematic consideration of how such hypothetical design space might be explored is beyond the scope of this study, this test case nicely illustrates how such an extension of GCAD may be motivated and even guided.

Another key lesson is the essentiality of explicitly considering drivers of cell-to-cell variation to obtain effective genetic program designs. In the framework used in this study, the primary driver of intercellular variation was variable uptake of plasmids. However, for any DNA delivery method (ranging from lentiviral and transposon vectors, which integrate pseudo-randomly in the genome, to CRISPR-mediated insertion of DNA in a single targeted locus), intercellular variation will still exist and may impact circuit performance. A good quantitative model of such variation is needed to incorporate this consideration into circuit design. Our Pulse Generator case study illustrated that explicitly optimizing for robustness to intercellular variation (e.g., maximizing the fraction of cells that will perform as desired despite intercellular variation) is a useful and at least somewhat feasible strategy. As in the Signal Conditioner case, any limited success with optimizing for robustness to variation could motivate prospective exploration of alternative design spaces, such as contemplating which DNA delivery method (and associated descriptors of intercellular variation) best enables meeting a design goal with a given set of parts.

Although we validated that GCAD can successfully recover optimal solutions, it is interesting to examine the limits of our method for this application. For the multi-objective cases, Signal Conditioner and Pulse Generator, our method utilized the non-dominated sorting procedure of NSGA II.^36^ While NSGA II is one of the most successful multi-objective evolutionary algorithms,^43^ it still faces limitations in approximating the full Pareto set as a result of local Pareto optimal solutions and biases induced by its crowding distance.^44^ This, in turn, limits our method’s ability to consistently recover the entire set of global Pareto optimal solutions. However, evolutionary-based algorithms are generally designed to approximate the set of Pareto optimal solutions as finding the exact Pareto set is expensive and usually infeasible.^45^ From this perspective, GCAD has a relatively strong performance since it can recover most of the Pareto optimal solutions, and thus any limitations appear unlikely to prevent selection of multiple optimal or near-optimal solutions. Nonetheless, it would be informative to investigate whether multi-objective searches are improved by experimenting with different sorting algorithms that tackle the limitations of NSGA II.^44, 46^

Finally, this study motivates several potential future directions for mammalian genetic circuit design automation employing the GCAD framework. First, the existing framework could be adapted to expand the design space by allowing for more parts in each circuit topology, even while using only the existing set of parts. In this study, we set the maximum number of parts in each circuit to 2 and placed constraints on topology (**Methods**) to focus on GCAD method development. However, simply enabling GCAD to search for more complex circuits designs could yield improved performance (e.g., enable design of Signal Conditioner circuits) or could generally enable design of circuits that can achieve more complex functions. Scaling the design space would result in additional computational expense associated with each GA run, but existing parallel genetic algorithm methods^47^ could be employed to address this challenge. Second, GCAD could be modified to enable design with an expanded or different set of biological parts. As discussed above, the addition of new parts would necessitate development of new (and ideally high confidence) models to guide utilization of these parts, and exploration of hypothetical design space could help to focus experimental work to develop new parts. Third, explicitly incorporating ODE model uncertainty into GCAD searches could prioritize solutions that are less subject to model uncertainty, or simply reporting model-prediction uncertainty would provide users with guidance about lower confidence designs. Evolution guided Bayesian optimization has recently been used in self-driving labs to accelerate the multi-objective optimization of materials.^48^ It is interesting that within our case studies, the ability to select useful topologies may have been less impacted by such model uncertainty, as many optimal (or Pareto optimal) circuits shared a topology but differed in parameters associated with part choice. Overall, the GCAD framework comprises an advance in semi-automated design that has the potential to accelerate the design process for building useful, high-performing mammalian genetic programs for diverse applications.

## Methods

### Developing a genetic parts model

Although our model is based on a published model for describing COMET synTFs and promoters,^35^ we made several key modifications for the purposes of this study.

#### Simplifying the representation of population-level heterogeneity

We used a reduced version of the representation of population-level heterogeneity to increase computational efficiency in GA runs. The original model employed a representation of population-level heterogeneity due to differences in DNA plasmid uptake across transiently transfected cells.^35^ A Gaussian mixture model was fit to the experimentally observed distribution of DNA uptake across a cell population, and samples were drawn from this distribution to generate a simulated population of cells, each with a different amount of DNA plasmid uptake for each plasmid that comprises the genetic circuit (i.e., within a single cell, each plasmid was individually assigned a DNA uptake value such that the ratios of plasmids of different types taken up can vary between cells). For a given circuit, a simulation was run for each cell in the simulated population, from which the average population-level behavior was calculated.^35^ The original model employed a 200-cell population, but the computational expense associated with running 200 simulations for each circuit in the search space made this model size intractable for implementation. We reduced the model to a 20-cell population, which enabled reasonable simulation during the GA. For each test case, we compared simulated, population-level average optimization objectives between the 20 and 200-cell models to ensure that the 20-cell model could sufficiently recapitulate behavior seen with the 200-cell model for the optimization objectives use in GCAD (**Supplementary** Figures 3**, 4, 10** and **Figure 4c**).

#### Adapting the promoter activation function

We generalized the functions for promoter activation in the genetic parts model based on hypothesized behavior of regulatory interactions to enable generation of ODEs for all of the circuits in the GCAD design space. The original model used the sigmoid function below to describe activator-dependent promoter activation (i.e., how much DNA is transcribed based on the amount of activator and repressor present).^35^

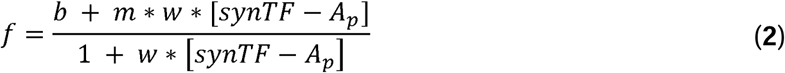

Here, the parameters (*b, m,* and *w*) represent the leaky promoter activation in the absence of synTF-A, the maximum promoter activation, and the steepness of the activation function, respectively, for the given synTF-A variant. **Table 2** lists the kinetic parameters for the synTF variants used here.

**Table 2.** Promoter activation parameters for synTFs.

| synTF variant | $b$ | $m$ | $w$ |
| --- | --- | --- | --- |
| 1 | 0.08 | 33 | 0.036 |
| 2 | 0.25 | 54 | 0.018 |
| 6 | 0.02 | 58 | 0.043 |
| 7 | 0.11 | 46 | 0.025 |
| 8 | 0.07 | 43 | 0.041 |
| 9 | 0.46 | 33 | 0.096 |
| 10 | 0.01 | 31 | 0.037 |
| 11 | 0.08 | 32 | 0.025 |
| 12 | 0.15 | 33 | 0.065 |
| 13 | 0.04 | 41 | 0.012 |
| 14 | 0.2 | 30 | 0.069 |
| 15 | 0.18 | 33 | 0.007 |

For cases with activation and repression with the repressors used here, the previously developed model used a promoter activation function (**Equation 3**). However, the model only described circuits in which the activator and repressor regulating a reporter protein were the same variant. Later work extended the model representation to describe the case in which the activator synTF1-A and repressor synTF2-R regulate a reporter, which we used as a basis for generalizing the promoter activation function^6^. We represented simultaneous activation and repression by any synTF variant pair using a similar formulation to that described by Muldoon et

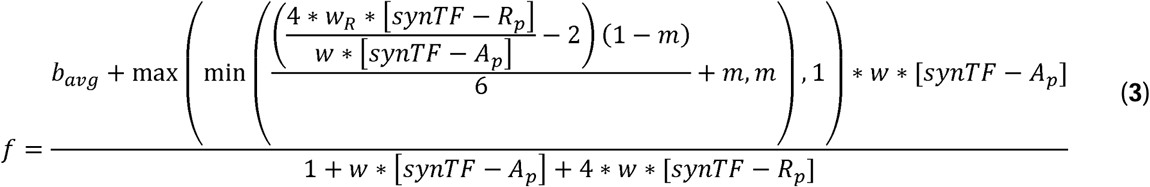

Here, *w_R_*is the *w* value for the synTF-R variant, and *b_avg_* is the average *b* value of the two synTF variants, which differs from the assumption used by Muldoon et al for the *b* value.^6^ The parameters *b*, *m*, and *w* here are the same as those in **Equation 2**.

Because cases with dual activation by different activator variants were also not directly empirically characterized previously, we used the following, hypothesized functional form for the dual activator regulation:

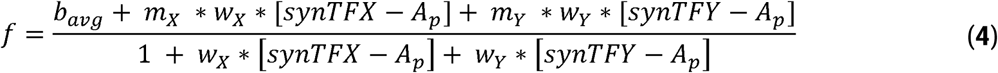

Here, synTFX-A and synTFY-A each represent a synTF-A variant, and *m_X_, m_Y_,* etc. indicate the variant-specific parameters (**Table 2**). As in **Equation 3**, *b_avg_* is the average of the *b* for the two synTF variants.

The dynamics of any circuit can then be described by the following system of ODEs:

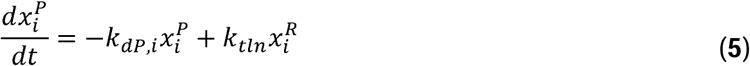

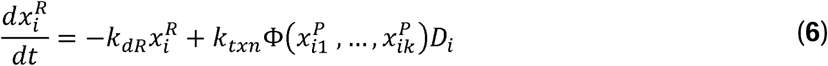

x^P^_i_ and x^R^_i_ are the concentration of mRNA and protein for component I, which can be an activator, repressor or the reporter. Note that promoters are not included in the system because they are not represented as dynamic quantities. k_dR_, k_dP,i_, k_txn_, and k_tln_ are fixed parameters describing the mRNA degradation rate, protein degradation rate, transcription rate, and translation rate, respectively, for each component, and are defined in **Table 3**. If the component is a synTF, the protein degradation rate is k_dP,syn_, whereas if the component is the reporter, the protein degradation rate is k_dP,rep_ D_i_ is the dose of DNA plasmid and Φ(x^P^_i1_, …, x^P^_i1_) is a generalized promoter activation function that takes the form of **Equation 2**, **3**, or **4**, depending on the regulation of the component within the circuit.

**Table 3.**
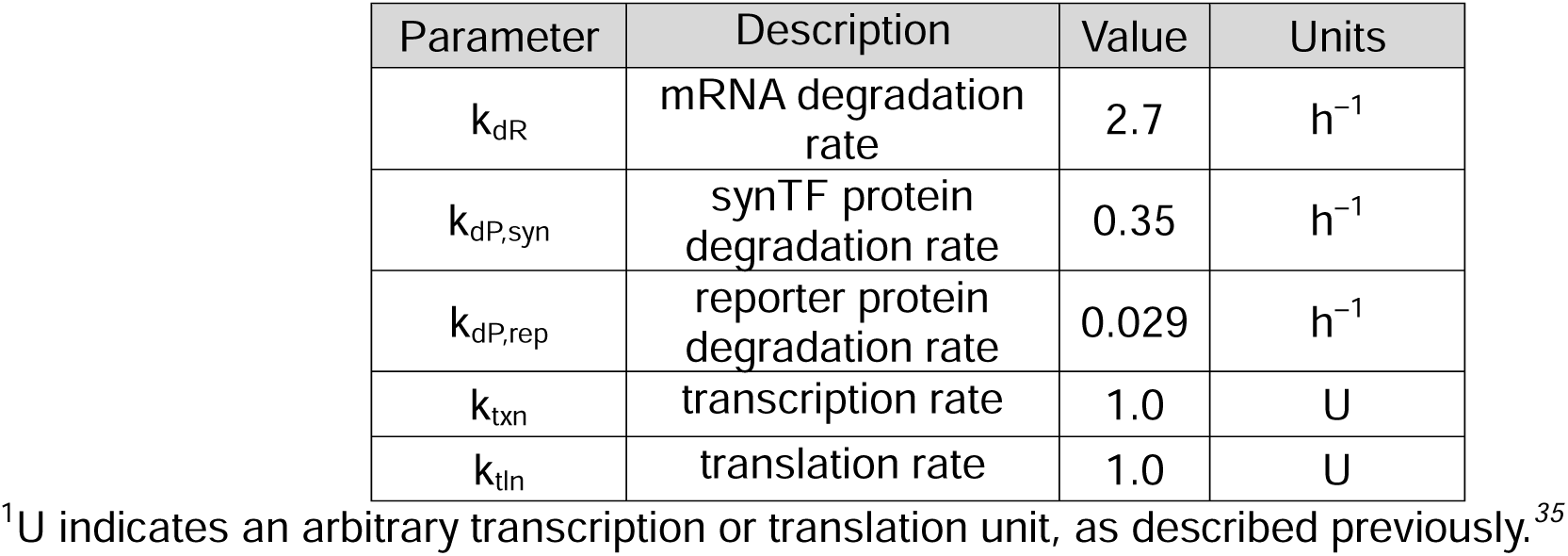
Fixed transcription, translation, and degradation parameters. ^1^

### Genetic algorithm for circuit selection

Genetic algorithm (GA) is an ensemble method that evolves a population candidate solution over a number of generations; only the “fittest” candidates survive and advance from one generation to the next. GA works well for optimization problems with nondifferentiable objectives and/or discrete variables, and it is more computationally efficient than combinatorial search. GA also enables flexible update rules for the candidates, which can be designed to directly parallel typical biological mutations. Furthermore, extensive benchmarking studies in the field of chemical engineering have shown that GA consistently outperforms more complex neural network-based methods in designing chemical molecules^49, 50^ and should be considered an important baseline. Given its strengths and merits, GA is well-suited for the automatic design of genetic circuits. (See **Supplementary Note 1** for how our method differs from previous works with GA).

#### Overall workflow

The GA procedure is as follows. We start by sampling a population of N candidate graphs. To account for the different combinatorial space of graphs of varying sizes, we can incorporate biases in sampling graphs with larger number of nodes. We next select parents using tournament selection and perform crossover with a set probability; this process is repeated until we obtain N children. For each child, we also perform mutation with a set probability. We tested the performance of GCAD with varying crossover and mutation probabilities to assess performance and determine baseline values (**Supplementary Note 2**, **Supplementary** Figures 16-18). Repair is applied after crossover and after mutation. The parents and children are then ranked based on their fitness values. The top N performing graphs are retained and form the pool of parent candidates for the next generation.

It is worth noting that crossover for graphs introduces additional mutation in the process regardless of how the parents’ fragments are split or recombined because biological constraints prevent a direct cut-and-paste procedure like with string representation. Moreover, the repair step can be conceptualized as another layer of mutation. As a result, unlike existing GA methods in which a topology undergoes more than one kind of mutation, we take a more restrained approach applying only one mutation at a time. In practice it is difficult to determine what the right combination of mutations should be and whether a combination found for one problem would work for a wide variety of problems and objective functions. Thus, if a graph is selected to mutate, we randomly use one of the four operators, each with equal probability.

#### Graph-based representation of circuits

Here, we encode each circuit as a directed graph in which each node in the graph represents a component in the genetic parts library, and each edge indicates a regulatory relationship. By default, the endogenous promoter node and the reporter node are present in every circuit. GA evolves the number of activator and repressor nodes, the number of edges in the circuit, the activator and repressor variant, and the plasmid dose of each component. An edge and its direction represent the regulation among the nodes. If there is an edge from node A to node B, then A regulates the production of B. Node A is also referred to as a predecessor of node B and node B as a successor of node A. The edge pointing from A to B is also known as an in-edge for B and as an out-edge for A. Additionally, components can self-regulate, captured by a loop edge that connects the node to itself.

#### Invalid and redundant topologies

The biological properties of genetic components put restrictions on the graph configurations that are valid. **Figure 5** shows examples of a valid graph (left panel) and invalid or redundant graphs (right panel). Invalid topologies are those that are not biologically functional. An example is a graph in which there is no activator node (**Figure 5b**) or a graph in which an activator only has an in-edge from itself and not from the promoter or other activators (**Figure 5c**). This means that there is no protein production for the reporter, which renders the circuit non-functional. Another example is a graph in which an activator and its corresponding inhibitor are both in the circuit, but the nodes do not have the same set of successors (**Figure 5d**). The invalidity of this graph is library dependent. On the other hand, redundant topologies are those instances that effectively result in the same dynamics for the reporter as another, often simpler, topology. For example, if a node has no path from itself to the reporter as in (**Figure 5e**), it has no effect on the reporter protein. Additional restrictions based on biological function or desired properties could be incorporated into the algorithm. Here, we chose a minimal set which would produce activation of the reporter from the promoter, maintain consistency in part identity, and avoid tracking parts which have no influence on the dynamics.

**Figure 5.**
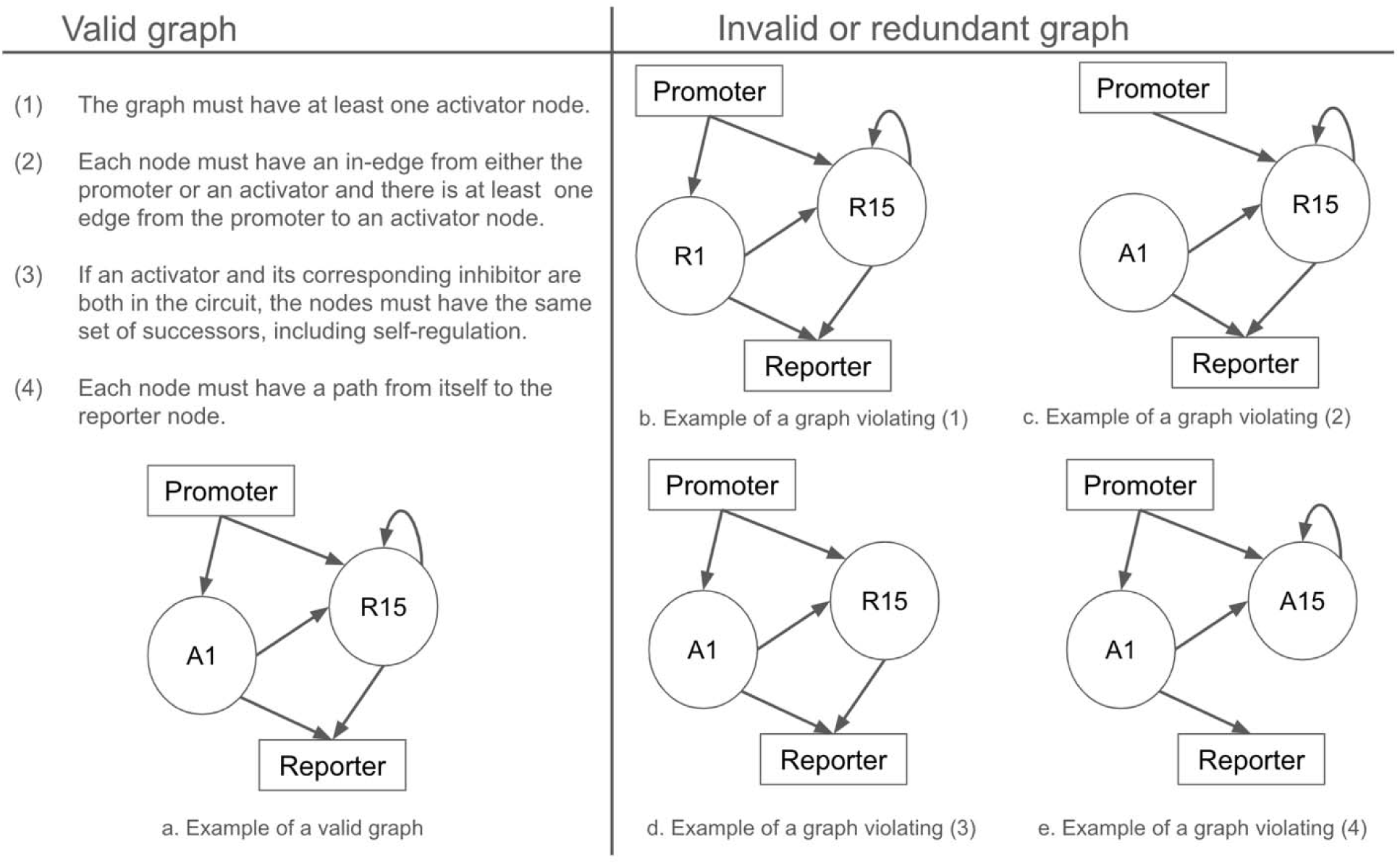
Rules to enforce validity of a circuit topology with example of a valid graph (a) and invalid (b, c, d) or redundant (e) graphs.

#### Crossover

Crossover is a fundamental operator of GA and previous work demonstrated strategies for crossover within graph-based encodings. Globus et al. formulate a crossover procedure for chemical molecules as undirected graphs.^51^ The operator first removes a set of edges whose deletion results in two disconnected fragments. The operator then matches the edge-cut-set between one parent’s fragment with the other’s. A cut edge is selected randomly from one fragment. If there exists an edge in the other parent’s fragment of the same bond type, the nodes from these fragments are joined with such bond type. If no compatible edge exists, a random edge is selected, and the nodes are connected with a new edge that satisfies valence.

As pointed out by Brown et al., this operator, termed multipoint crossover, can break cycles in the graph and it is not possible to control the size of the fragments, and consequently the resultant children.^52^ To mitigate this issue, they propose a subgraph crossover. The operator derives a connected subgraph from each parent and joins these two subgraphs in the same manner as by multipoint crossover. The creation of a connected subgraph entails recursively selecting an edge that is incidental with any previously chosen edges. This stops when the subgraph is roughly half the size of the parent graph.

For our work, we propose a node-based crossover that draws inspiration from the point-based crossover of binary string of the classical GA. The operator can also be conceptualized as a subgraph crossover in which the subgraph is created by randomly choosing a node along with its edges. This is also consistent with the biological interpretation that each node is a regulatory element which can be selected upon and who’s connections can be modified during the GA process. To obtain the parent candidates, we use tournament selection.^53^

#### Node-based crossover

The node-based operator first identifies a crossover node for each parent. We prioritize nodes that are common in both parents. If there are no overlapping nodes, we first pick a node from one parent and pick a node of the same type (activator or inhibitor) from the other parent. If one of the parents does not have any inhibitor nodes, we pick from the available nodes.

After determining crossover nodes, we remove the selected node along with its in and out-edges from the parent. This essentially breaks the parent graph up into two fragments, *F_1_*, containing the crossover node and its edges and *F_2_*, the remaining portions of the graph. The procedure then joins the *F_2_* fragment of one parent with the *F_1_* fragment of the other. The goal is to retain as much of the in and out-edges of the crossover node along with the type of its successors and predecessors in the new graph as possible. Biologically this means that as much local regulatory structure is maintained as possible.

**Figure 6.**
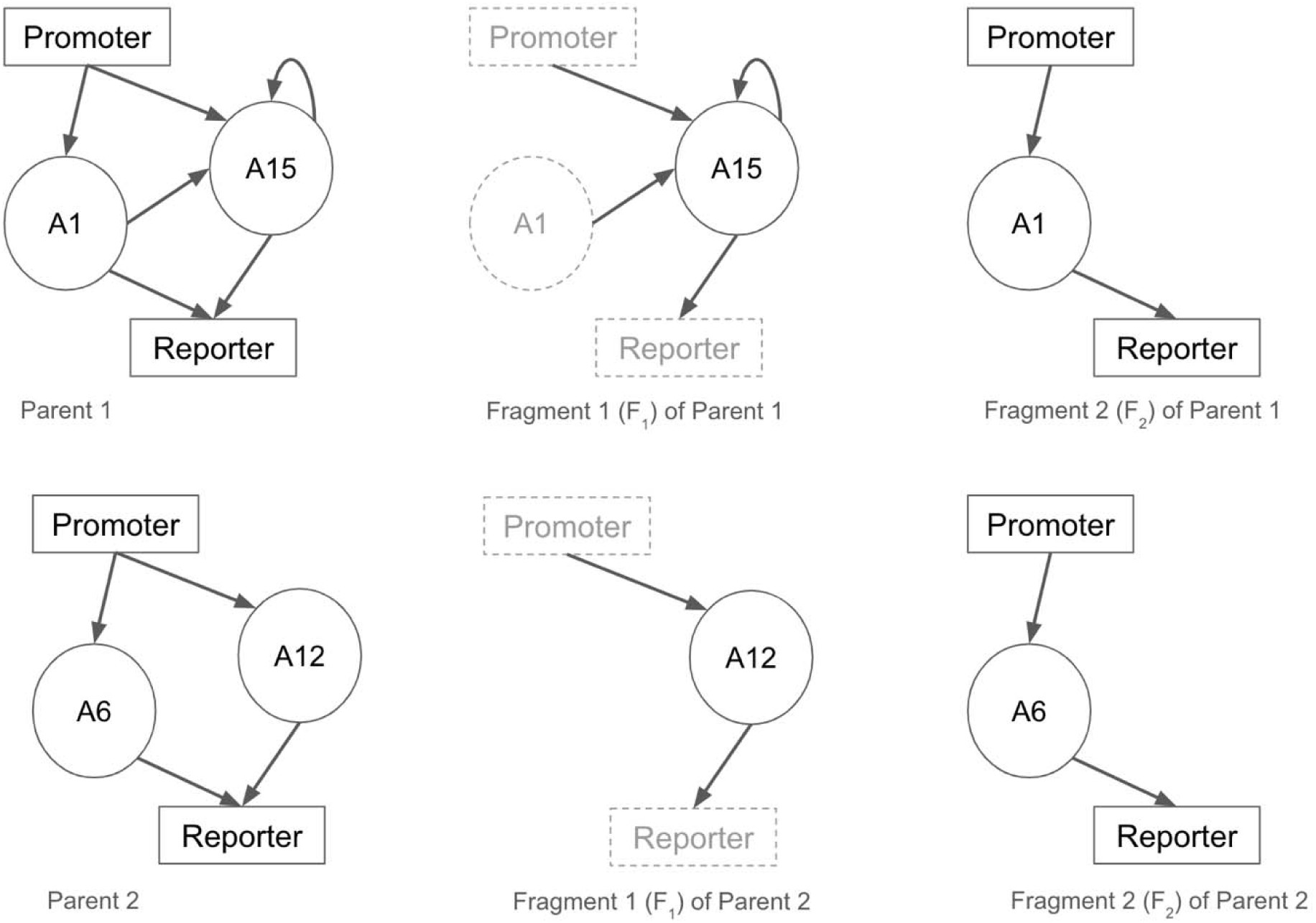
Splitting parent graphs into fragments based on crossover nodes.

There are two key rules for the crossover operator. First, the crossover node’s loop edge, its in-edge from the promoter node, and its out-edge to the reporter are always retained in the children graph. Second, if the node’s successors and predecessors from the parent graph are already present in the new *F_2_* fragment, the edges are kept between the same nodes in the new graph. Otherwise, we randomly pick a node of the same type as that from the parent in the new *F_2_* fragment and create a new edge.

**Figure 7.**
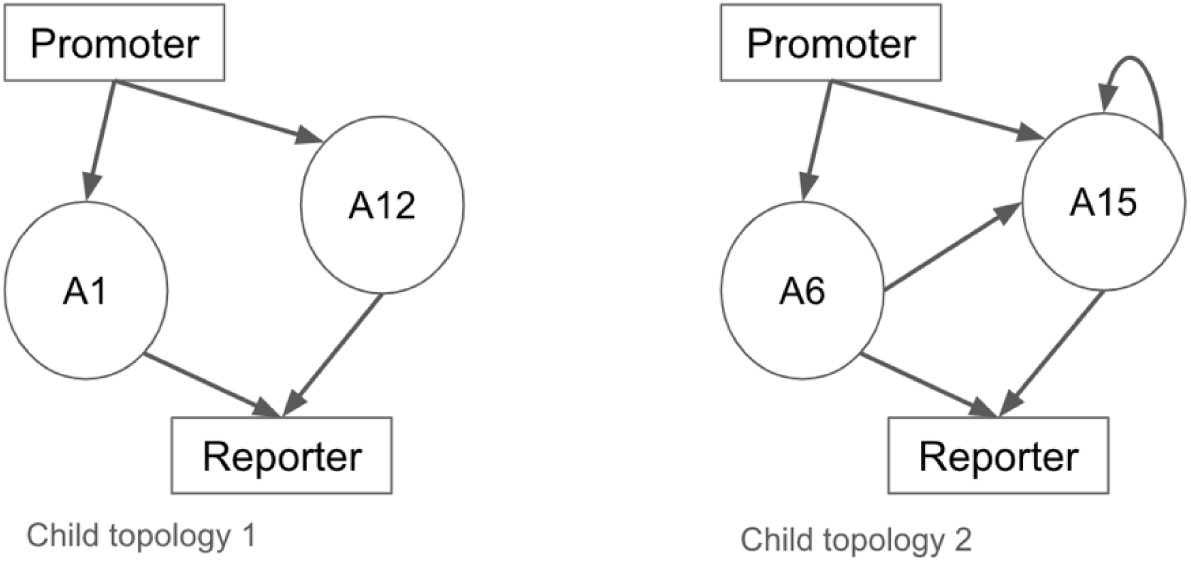
Children graphs resulting from crossover of parent graphs in Figure 2.

#### Mutation

A graph-based representation includes four key pieces of information: the number of activator and inhibitor nodes, node identities from the parts library, the in/out edges that embed regulatory relationship, and the plasmid doses. We adapt the possible mutation operators to target each of these properties. Note that these mutations are similar to the procedures proposed by Brown et al.,^52^ except for the plasmid doses.

#### Mutation 1: number of activator/inhibitor nodes

We can either insert a new node into the graph or delete an existing one. To insert a node, we pick a node from the library that is not currently used and insert it into the graph by randomly creating edges to other nodes such that the resultant graph is valid. To delete a node, we select an existing node such that without it, the remaining graph will still have at least one activator node.

If a graph has only one node in addition to the promoter and reporter nodes, we implement the insertion operator; if a graph has the maximum number of nodes as prescribed by the user, we implement the deletion operator. Otherwise, we randomly choose from the two operators with probability *p* = 0.5.

#### Mutation 2: node identity

In addition to the number of activators and inhibitors, we can select a different node of the same type from the library to use in a circuit. If the node corresponds to an activator (inhibitor), the operator replaces it with a different activator (inhibitor) in the library that is not already present in the graph. The corresponding plasmid dose and the existing in / out edges are retained.

#### Mutation 3: edge mutation

Similar to modifying the number of nodes, we mutate edges through either insertion or deletion. Regarding insertion, to ensure that the circuit is consistent with the constraints of the genetic parts library, new out-edges from inhibitor nodes must be treated carefully. Specifically, if an inhibitor is selected for edge insertion and its corresponding activator is present in the circuit, the new out-edge is added to both nodes to ensure they have the same set of successors. Removing a randomly selected edge can frequently render a graph invalid. Therefore, we randomly select an edge and check that its removal will not affect the validity of the resultant graph. If it does, we pick a different edge.

If all activator/inhibitor nodes are fully connected and properly connected to the promoter and the reporter, we implement the deletion operator. If the graph is valid and each node has exactly one in-edge and one out-edge, we perform insertion. Otherwise, we choose one of the two operators with a probability *p* = 0.5.

#### Mutation 4: dose

This mutation operator perturbs the dose of the corresponding genetic component of a node. In this paper, doses are in the range of 5 to 75 (inclusive), and to mutate the dose of a node, we randomly draw an integer multiple of 5 in this range.

#### Other operators

Besides crossover and mutation, the sampling of the initial population also requires special consideration for graph encoding. To generate a circuit topology, we first determine the desired number of activators/inhibitors and randomly select the nodes from the library. After that, we construct a set of incidental edges that form a path from the promoter to the node and another set that forms a path from the node itself to the reporter.

Repair is another operator that we adapt for graph representation of circuits. Both crossover and mutation can cause the resultant child topology to become invalid (e.g., an activator without any in-edges from the promoter or other activators) or redundant (e.g., a node having no path from itself to the reporter (**Figure 5e**) and, thus no effect on the reporter protein). While it is possible to remove these topologies and create new ones, that can be computationally wasteful and potentially lead to a time-consuming loop. Instead, we check the validity of new topologies according to the four rules listed in **Figure 5** and employ the repair operator. Note that in designing crossover and mutation, we are careful to maintain the node identity, thus ensuring the validity of the set of nodes. Therefore, we only need to check if a topology has a valid set of edges. This means determining if each activator/inhibitor node has at least a path from the promoter and a path to the reporter and if an activator and its corresponding inhibitor in the topology have the same set of successors. If not, we repair the graph by generating paths that meet these conditions. Otherwise, no repair is needed.

#### Hyperparameter optimization

We used response surface methodology^39, 40^ (RSM) to optimize genetic algorithm hyperparameters (**Figure 3c**). RSM is an approach for solving multi-objective optimization problems that has previously been utilized for hyperparameter optimization of GAs^54^ and other machine learning algorithms,^55^ and it is particularly useful when running the simulation is computationally expensive. RSM involves fitting a simplified, surrogate function to each of the optimization objectives to bypass the need to run the full simulation to evaluate a set of candidate hyperparameters. The approach begins with a global search using semi-random, Latin hypercube sampling^56^ of the hyperparameter space to generate simulation data—in this case, from running the GA. We used a sample size of 200 hyperparameter sets for this step, and the search space consisted of probabilities of crossover and mutation between 0 and 1; we chose to fix the size of the population used to initialize the GA at 200 and the ratio of 1 to 2-part circuits in the population to 0.25. Next, the global search simulation data is used to fit a surrogate function—here, we used a Gaussian radial basis function—to each of the optimization objectives, which maps the hyperparameter space to the objective space. For the GA hyperparameter optimization, we chose to maximize the hypervolume of the solution set and minimize the number of generations to convergence, i.e., the first generation at which the final hypervolume is obtained. The set of surrogate functions is then transformed into a group of n_scalar_ single objective subproblems (we used 20) via scalarization, e.g., a weighted average of the objective values, for optimization. Each of the n_scalar_ subproblems is then optimized to generate n_scalar_ candidate solutions (hyperparameter sets), and the GA is evaluated for each candidate solution. The evaluation data is used to refit the set of surrogate functions, and the process is repeated until the termination criteria is met, at which point the non-dominated hyperparameter solution set is returned. We used the number of surrogate function fitting iterations as the termination criterion; we set this termination criterion to 10 iterations. We utilized the parMOO algorithm^40^ and associated Python package^57^ to perform the RSM optimization.

#### Representing inducible constructs for reference circuits and endogenous promoter proxies

As described in the main text, a dox dose-response experiment was used to select dox concentrations to represent the OFF and ON states of pEnd (**Supplementary Figure 6**). For subsequent experiments, the reference circuit was activated using the dox concentration corresponding to the OFF and ON states of pEnd. For describing these parts in the model, we first calculated an effective transcription rate for the OFF and ON state using an analogous method to the one used to calculate the *m* parameter in the previously developed, genetic parts model,^35^ We then incorporated the pEnd parameters into the model ODEs for parts regulated by pEnd.

### Experimental methods

#### General DNA assembly

Plasmid cloning was performed primarily using standard oligo annealing and restriction enzyme cloning with Phusion DNA Polymerase, restriction enzymes (NEB; Thermo Fisher), T4 DNA Ligase (NEB), Antarctic Phosphatase (NEB), and T4 PNK (NEB). Non-reporter plasmids were all created by restriction enzyme cloning consisting of one destination plasmid and one insert plasmid. Golden Gate assembly was also utilized to assemble reporter constructs. All plasmids were transformed into chemically competent TOP10 *E. coli* (Thermo Fisher) and grown at 37°C.

#### Source vectors for DNA assembly

Most parts were sourced from plasmids previously described but several were created in this study (**Supplementary Tables 1-3).** ZF binding sequences and synTF encoding sequences were taken from previous reporting of the genetic parts toolkit.^35^ EYFP was sourced from plasmids we previously described (Addgene plasmid #58855).^35^ EBFP2 was sourced from pEBFP2-Nuc, which was a gift from Robert Campbell (Addgene plasmid #14893). Plasmid TetON3G were sourced from pLVX-Tet3G (Clontech), and TRE3GV was sourced from pLVX-TRE3G (Clontech). All plasmid backbones are modified versions of the pcDNA3.1/Hygro(+) Mammalian Expression Vector (Thermo Fisher). <u>V87020</u>).

#### Golden Gate assembly of ZF reporter plasmids

Golden Gate assembly was used to construct the reporter plasmids from a reporter template (pD540). Promoter inserts were synthesized as 15–100 bp oligonucleotides (some promoters were synthesized as multiple inserts) by Integrated DNA Technologies or Life Technologies (Thermo Fisher). The coding and reverse strands were synthesized separately and designed to anneal, resulting in dsDNA with a 4Lnt sticky end overhang on each side. The coding and reverse oligonucleotides were mixed (6.5 µL H_2_O, 1 µL T4 Ligase Buffer, 0.5 µL T4 PNK (10 U/µL; NEB), 1 µL of each 100 µM oligonucleotide) and phosphorylated at 37 °C for 1 h. They were then denatured at 95 °C for 5 min and cooled slowly to room temperature (here, approximately 22 °C) to allow for annealing. The mix was then diluted 500-fold to make a 20 nM stock. These annealed oligos in conjunction with a reporter plasmid template vector were then used in a BsaI Golden Gate reaction to assemble the new reporter plasmid. BsaI Golden Gate reaction mixtures comprise 1 µL T4 ligase buffer, 1 µL 10× BSA (1 mg/mL), 0.5 µL BsaI-HF (20 U/µL; NEB), 0.5 µL T4 Ligase (400 U/µL; NEB), 10Lfmol of vector, 1 µL of each insert (diluted to 200 nM or 20 nM), and water to 10 µL total volume. The reaction was incubated at 37 °C for 1 h, 55 °C for 15 min, and 80 °C for 20 min, and then cooled to room temperature. Up to 10 µL of reaction was immediately transformed into up to 50 µL of chemically competent Top10 *E. coli*. For reactions that did not yield many colonies on the first cloning attempt or did not produce colonies with the correct plasmids, the reaction conditions were changed to: 30 cycles of 37 °C for 1 min then 16 °C for 1 min, 55 °C for 15 min, 80 °C for 20 min, and cool to room temperature.

#### Plasmid preparation

TOP10 or NEB Stable *E. coli* were grown overnight in 50-100 mL of LB with Ampicillin and DNA was prepped using a ZymoPURE II Plasmid Midiprep Kit (Zymo #D4201) by following the manufacturer’s instructions.

#### Cell culture

The HEK293FT cell line was purchased from Thermo Fisher/Life Technologies (RRID: <u>CVCL_6911</u>) and was not further authenticated. Cells were cultured in DMEM (Gibco 31600-091) with 10% FBS (Gibco 16140-071), 6 mM L-glutamine (2 mM from Gibco 31600-091 and 4 mM from additional Gibco 25030-081), penicillin (100 U/μL), and streptomycin (100 μg/mL) (Gibco 15140122), in a 37 °C incubator with 5% CO_2_. Cells were subcultured at a 1:10 to 1:20 ratio every 3–4Ld using Trypsin-EDTA (Gibco 25300-054).

#### Transfection experiments

Experiments were conducted by transient transfection of HEK293FT cells using the calcium phosphate method. For transfection experiments, cells were plated at a minimum density of 1L×L10^5^ cells/well in a 24-well plate in 0.5 mL of DMEM, supplemented as described above. After about 24 h, by which time the cells had adhered to the plate and grown to about 50% confluency in the well, they were transfected via the calcium phosphate method. Plasmids for each experiment were mixed in H_2_O, and 2LM CaCl_2_ was added to a final concentration of 0.3LM CaCl_2_. This mixture was added dropwise to an equal-volume solution of 2× HEPES-Buffered Saline (280 mM NaCl, 0.05 M HEPES, 1.5LmM Na_2_HPO_4_) and gently pipetted up and down four times. After 4 min, the solution was mixed vigorously by pipetting ten times. 100 µL of this mixture was added dropwise to each well of the plated cells, and the plates were swirled gently. The next morning, the medium was aspirated and replaced with fresh medium. In some assays, fresh medium contained doxycycline resuspended in absolute ethanol. At 36– 48 h post-transfection and at least 24 h post-media change, cells were harvested for flow cytometry by washing with PBS pH 7.4 and using 0.05% Trypsin-EDTA (Thermo Fisher Scientific #25300120) for 5 min followed by quenching with medium. Cell suspensions were pipetted and added to 1 mL of FACS buffer (PBS pH 7.4, 2–5 mM EDTA, 0.1% BSA). Cells were spun at 150×g for 5 min, supernatant was decanted, and fresh FACS buffer was added. All experiments were performed in biological triplicate. For time point experiments, all cells were plated, transfected, and given new media at the same time, but the time in which they were harvested for flow cytometry varied slightly as indicated above. DNA dosages and plasmids used in all experiments are listed in **Supplementary Data 2.**

#### Flow cytometry

Flow cytometry was run on a BD LSR Fortessa Special Order Research Product (Robert H. Lurie Cancer Center Flow Cytometry Core and Single Cell Genomics Core). Lasers and filter sets used for data acquisition are listed in **Supplementary Table 4**. Approximately 3,000– 10,000 single, transfected cells were analyzed per sample in transfection experiments. Transfected cells were identified using a separate, single transfection control fluorescent protein (e.g., EBFP2). Samples were analyzed using FlowJo v10 software (FlowJo, LLC). Fluorescence data were compensated for spectral bleed-through and converted to absolute units using Spherotech UltraRainbow Calibration Particles (URCP). The HEK293FT cell population was identified by SSC-A versus FSC-A gating, and singlets were identified by FSC-A versus FSC-H gating. To distinguish transfected from non-transfected cells, a control sample of cells was generated by transfecting cells with a mass of pcDNA (empty vector) equivalent to the mass of DNA used in other samples in the experiment. For the single-cell subpopulation of the pcDNA-only sample, a gate was made to identify cells that were positive for the constitutive fluorescent protein(s) used as a transfection control in other samples, such that the gate included no more than 0.1% of the non-fluorescent cells. More details and an example workflow are presented in **Supplementary Figure 15**.

## Abbreviations

CI: Confidence interval
DBTL: Design, build, test, learn
Dox: Doxycycline
GA: Genetic algorithm
GCAD: Genetic program computer-aided design
NSGA: Non-dominated sorting genetic algorithm
ODE: Ordinary differential equation
pEnd: endogenous promoter
pSyn: synthetic promoter
RSM: Response surface methodology
SynTF: Synthetic transcription factor
SynTF-A: Synthetic transcription factor activator
SynTF-R: Synthetic transcription factor repressor

## Author contributions

K.S.D.: conceptualization, data curation, formal analysis, investigation, methodology, resources, software, visualization, writing – original draft, writing – review & editing.

A.V.N. conceptualization, data curation, formal analysis, investigation, methodology, resources, software, visualization, writing – original draft, writing – review & editing. G.G.B.: data curation, formal analysis, investigation, resources, visualization, writing – review & editing. L.E.R.: data curation, formal analysis, investigation, resources, visualization, writing – review & editing. H.I.E.: conceptualization, formal analysis, writing – review & editing. J.Z.: investigation, formal analysis, writing – review & editing. E.A.: investigation, writing – review & editing. K.E.D.: conceptualization, writing – review & editing. J.N.L.: conceptualization, funding acquisition, project administration, supervision, writing – review & editing. N.M.M.: conceptualization, funding acquisition, project administration, supervision, writing – review & editing. Manuscript was reviewed and edited by all authors prior to submission.

## Supporting information

**SupplementaryInformation.pdf**: additional figures and notes.

**SupplementaryData1.zip**: plasmid maps.

**SupplementaryData2.zip** source data.

Supporting computer code is available at the git repository: https://github.com/Mangan-Group/GraphGA

## Acknowledgements

This work was supported in part by the National Institute of Biomedical Imaging and Bioengineering of the NIH under award number 1R01EB026510 (J.N.L.), 2R01EB026510 (J.N.L. and N.M.M.); the National Science Foundation under award number DGE-1842165 (H.I.E.) and MCB-1745753 (J.N.L) and DGE-2021900 to the Northwestern University Synthesizing Biology Across Scales National Research Training program (G.G.B.); the Northwestern University Flow Cytometry Core Facility supported by Cancer Center Support Grant (NCI 5P30CA060553); and the NUSeq Core of the Northwestern Center for Genetic Medicine. H.I.E was supported in part by the National Institutes of Health Training Grant (T32GM008449) through Northwestern University’s Biotechnology Training Program. A.C. was supported in part by the Northwestern University Graduate School Cluster in Biotechnology, which is affiliated with the Biotechnology Training Program. K.S.D. and A.V.N. were supported by the Interdisciplinary Cluster on Predictive Science and Engineering Design (PS&ED) at Northwestern University. E.A. was supported in part by a grant from Northwestern University’s Office of Undergraduate Research. This research was supported in part through the computational resources and staff contributions provided for the Quest high performance computing facility at Northwestern University which is jointly supported by the Office of the Provost, the Office for Research, and Northwestern University Information Technology.

