## Supplementary Materials for "GCAD: a Computational Framework for Mammalian Genetic Program Computer-Aided Design"

**Contents:**

Supplementary Figures 1-18

Supplementary Notes 1-3

Supplementary Tables 1-4

References cited in this document

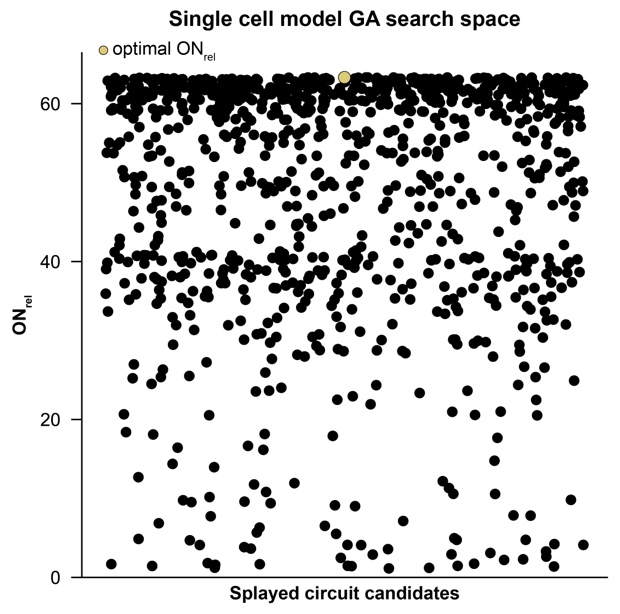

**Supplementary Figure 1. Single cell model GA search space for the Amplifier test case.** ON_rel_ values for the full GA search space with the single cell model are shown, and the optimal value from the combinatorial search is indicated. The GA search space spans the range of ON_rel_ from 0 to the optimal value, as was the case in the combinatorial search.

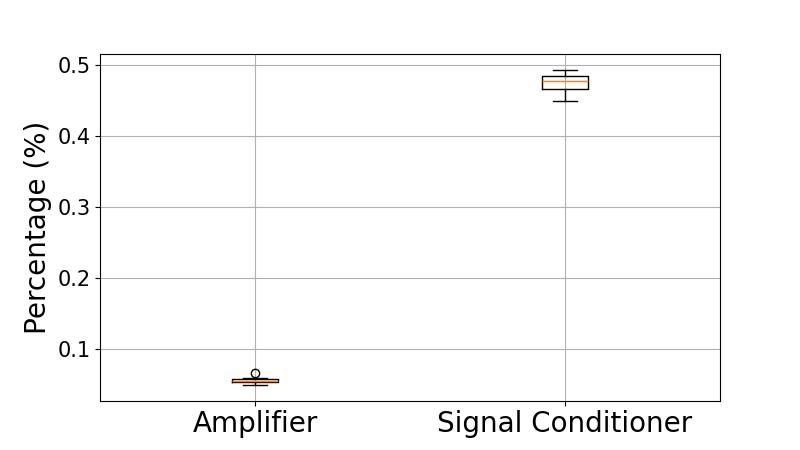

**Supplementary Figure 2. Percentage of unique topologies explored by GCAD over the total number of unique topologies found by combinatorial search.** Results for the Amplifier and the Signal Conditioner test case are obtained with the GA hyperparameters in **Table 1** after 120 generations. We differentiate topologies by different nodes, edges, and dose combinations, and there is a total of 1,679,760 topologies in the combinatorial search space.

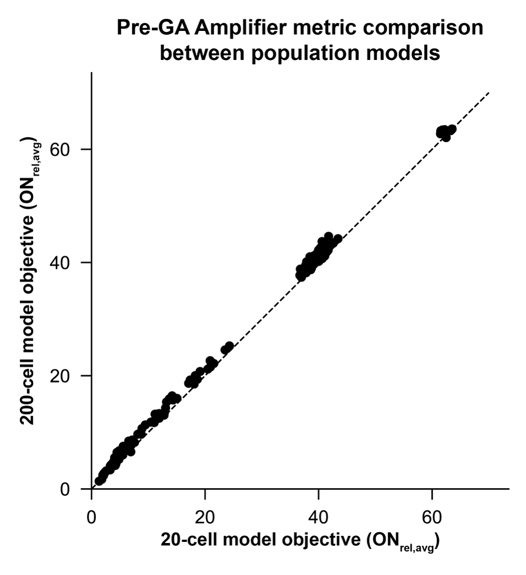

**Supplementary Figure 3. Pre-GA population model metric comparison for the Amplifier test case.** Selected circuits with low, mid-range, or high ON_rel_ values from the GA search with the single cell model were simulated with both the 20-cell population model and the 200-cell population model, and ON_rel,avg_ was calculated for each case. The dashed, y=x line indicates perfect agreement in ON_rel,avg_ between the two models.

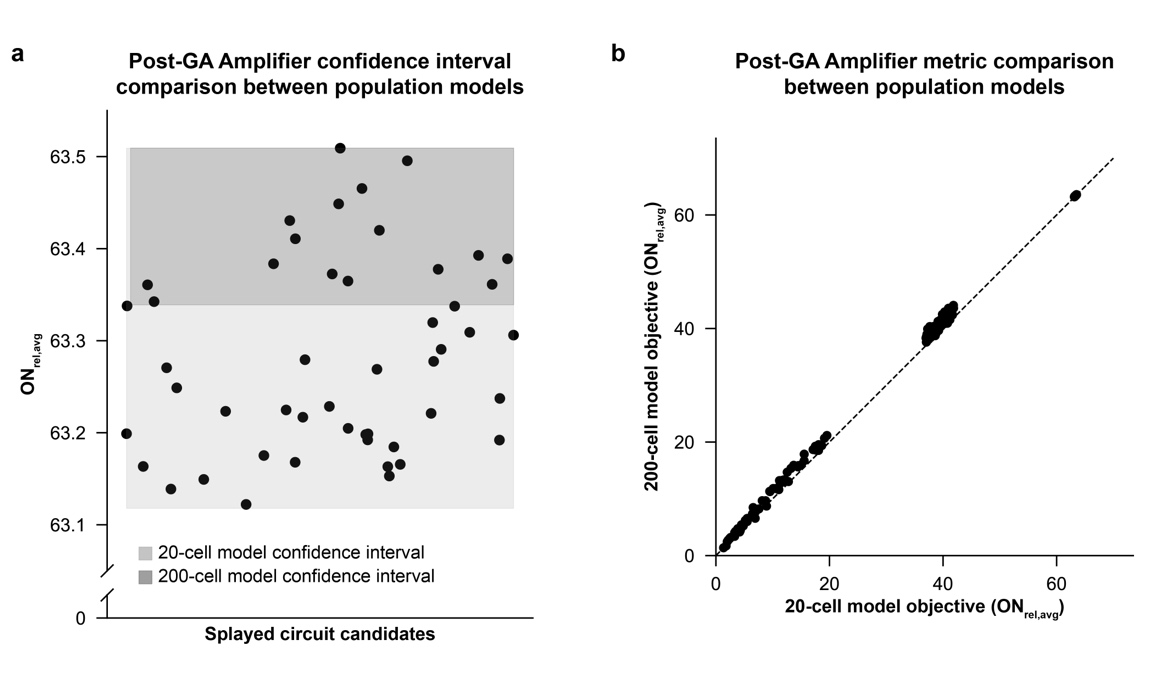

**Supplementary Figure 4. Post-GA population model metric comparison for the Amplifier test case.** (**a**) Circuits used to calculate the 20-cell confidence interval were simulated with the 200-cell model to calculate a 200-cell model confidence interval. The resulting confidence interval fell within the 20-cell confidence interval (overlapping with the top of the 20-cell confidence interval on the top of the plot). (**b**) Selected circuits with low ON_rel,avg_, mid-range ON_rel,avg_, or ON_rel,avg_ values within the confidence interval from the GA search with the 20-cell model were simulated with the 200-cell population model, and ON_rel,avg_ was calculated for comparison to the 20-cell model ON_rel,avg_ values. The dashed, y=x line indicates perfect agreement in ON_rel,avg_ between the two models.

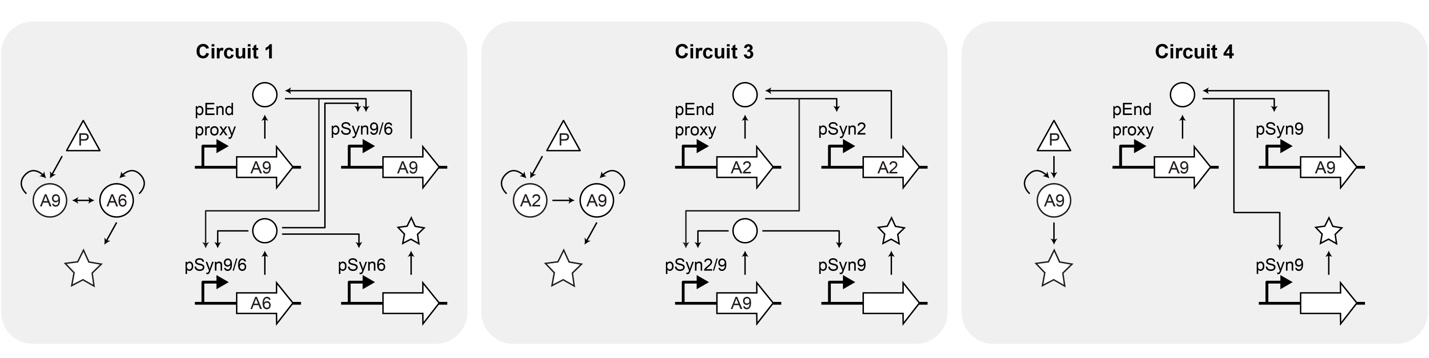

**Supplementary Figure 5. Amplifier experimental circuit directed graphs and circuit diagrams.** Selected amplifier topologies for experimental implementation translated from directed graphs to circuit diagrams. Circuit 2 is shown in **Figure 2e**.

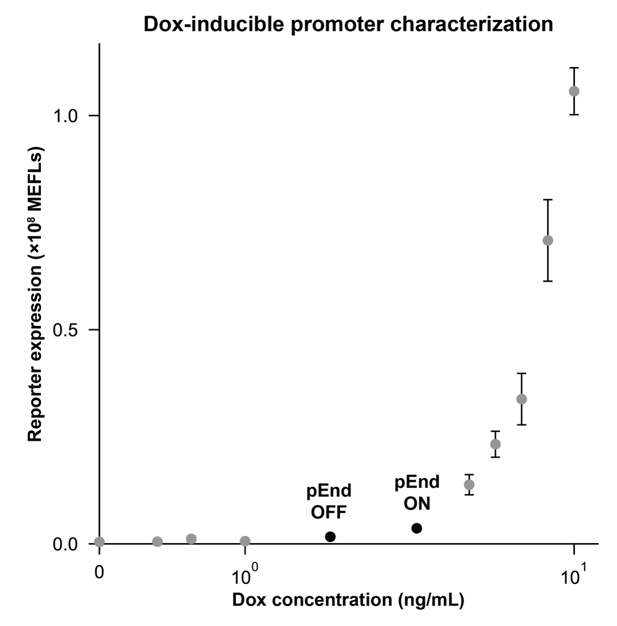

**Supplementary Figure 6. Characterization of endogenous promoter (pEnd) proxy.** A doxycycline (dox)-inducible Tet-On system was used as a proxy for pEnd. Tet-On-driven reporter expression was quantified at various dox concentrations, from which pEnd OFF and ON-state concentrations were selected; OFF = 1.58 ng/mL and ON = 2.51 ng/mL.

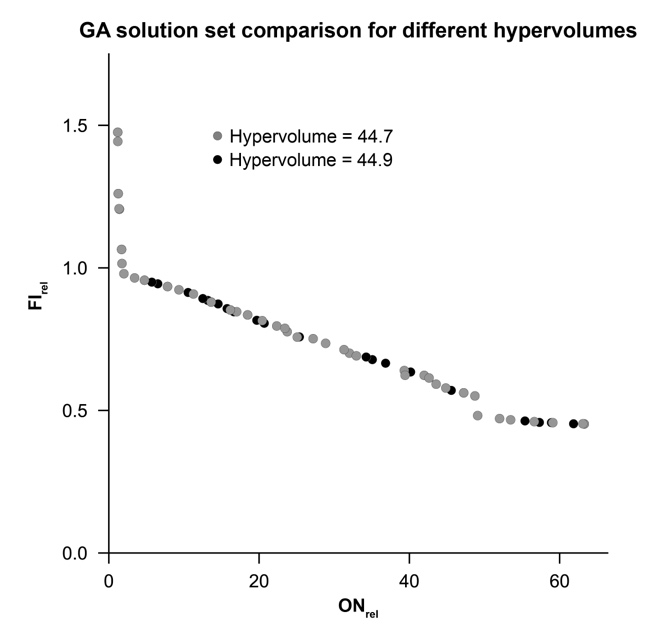

**Supplementary Figure 7. Comparison of solution sets between GA generations for the Signal Conditioner test case with initial hyperparameters.** The GA solution set at different generations within the same GA run using initial hyperparameters, with the indicated hypervolumes. The solution set with the lower hypervolume does not include several of the solutions in the higher hypervolume case due to repeated solutions, which results in a lower hypervolume.

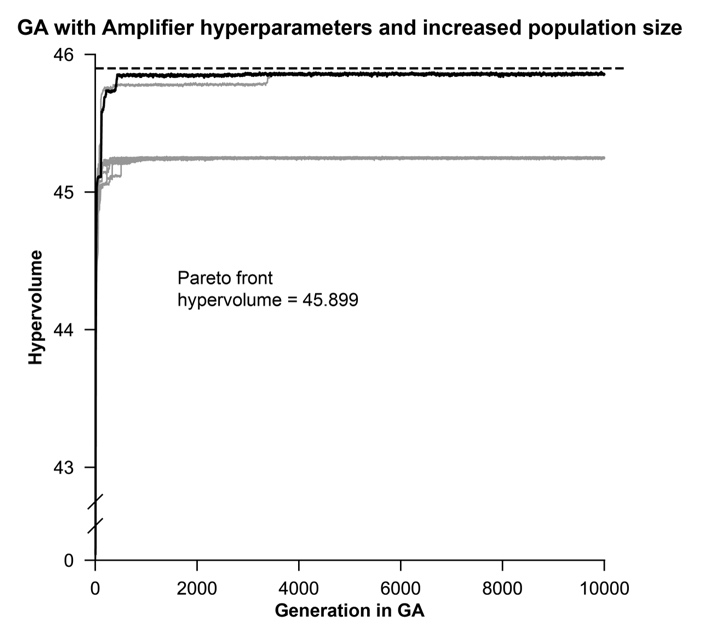

**Supplementary Figure 8. Hypervolume convergence for the Signal Conditioner with initial hyperparameters and increased population size.** The hypervolume converges near the Pareto front hypervolume (within 0.05) for 2/10 seeds, but the remaining seeds converge to a lower hypervolume (within 0.66 of the Pareto front hypervolume).

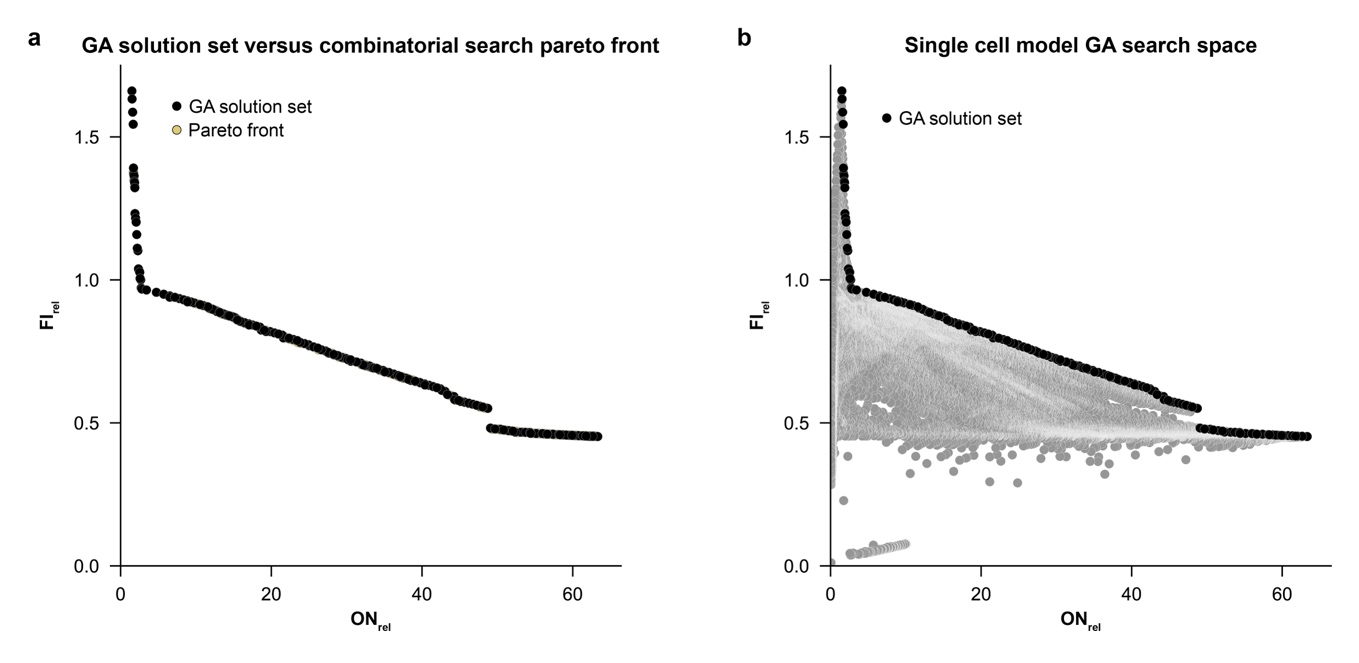

**Supplementary Figure 9. Single cell model GA search space for the Signal Conditioner test case.** (**a**) ON_rel_ and FI_rel_ values for the full GA search space with the single cell model are shown, and the nondominated solution set from the GA is indicated. The GA search space spans the range of ON_rel_ and FI_rel_ values from 0 to the maximum for each metric, with most points falling in the region with the most solutions indicated in the combinatorial search. (**b**) The GA solution set from (**a**) compared to the pareto front from the combinatorial search (the solution set overlaps with the pareto front, so it is not visible on the plot).

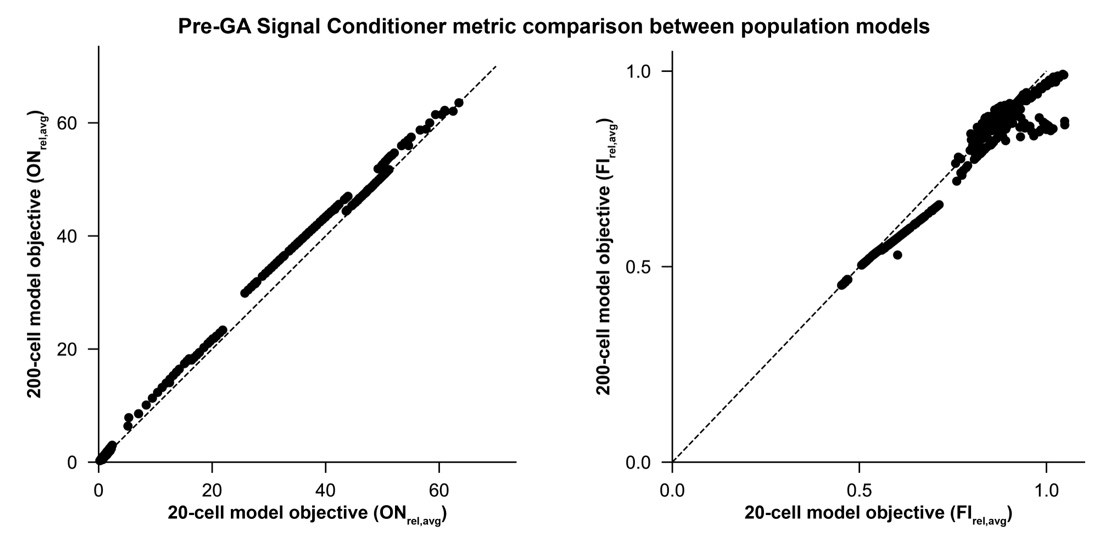

**Supplementary Figure 10. Pre-GA population model metric comparison for the Signal Conditioner test case.** Selected circuits with low ON_rel_ and FI_rel_, mid-range ON_rel_ and FI_rel_, or ON_rel_ and FI_rel_ values in the solution set from the GA search with the single cell model were simulated with both the 20-cell population model and the 200-cell population model, and ON_rel,avg_ (left) and FI_rel,avg_ (right) were calculated for each case. The dashed, y=x line indicates perfect agreement in metric between the two models.

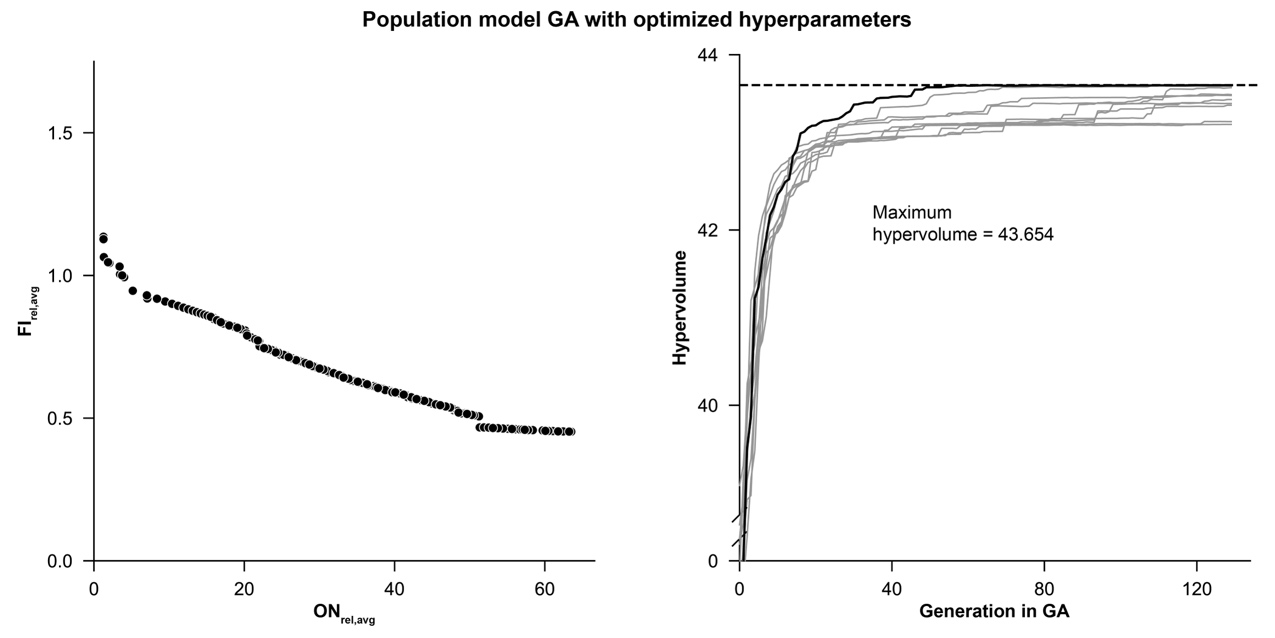

**Supplementary Figure 11. Model selection with the 20-cell model for the Signal Conditioner test case.** The solution set for the 20-cell model GA is shown for a selected seed with the optimized hyperparameters (left). The right plot shows the progression of the hypervolume through each generation in the 20-cell model GA for 10 different initialization seeds with the hyperparameters optimized for the single cell model (**Table 3**), with the maximum hypervolume indicated (dashed line).

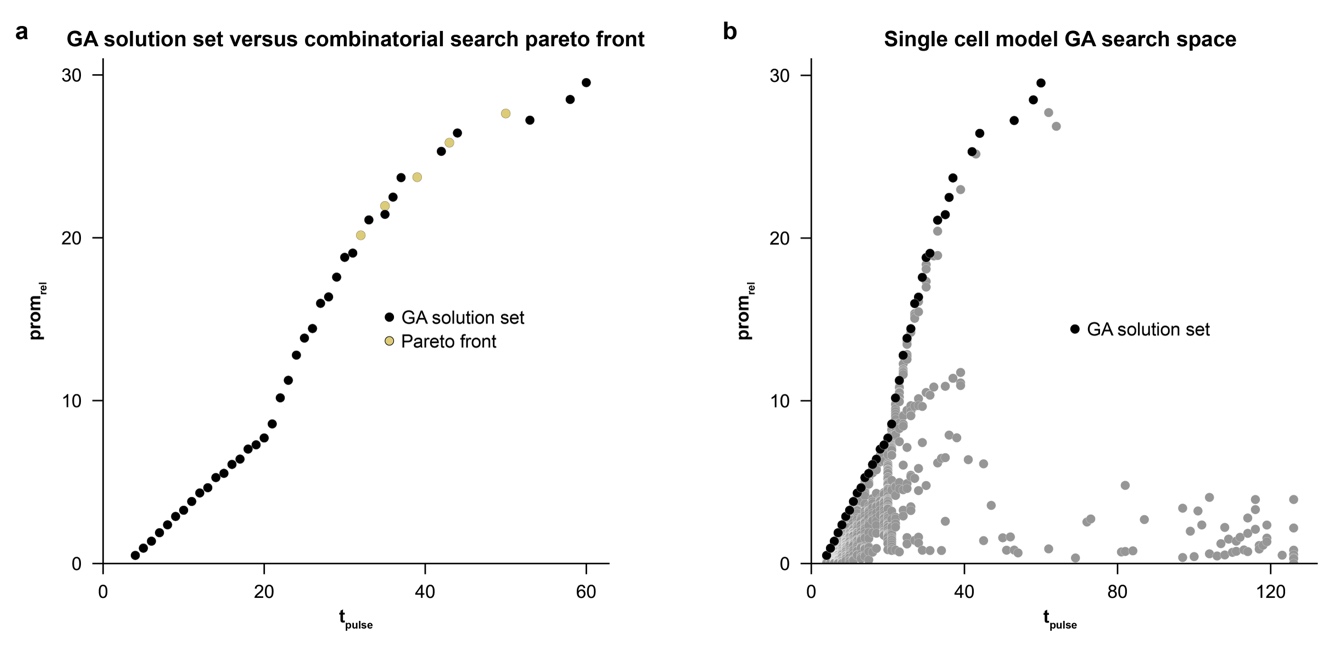

**Supplementary Figure 12. Single cell model GA search space for the Pulse Generator test case.** (**a**) t_pulse_ and prom_rel_ values for the full GA search space with the single cell model are shown, and the nondominated solution set from the GA is indicated. The GA search space spans the range of t_pulse_ and prom_rel_ values from 0 to the maximum for each metric, with most points falling in the region with the most solutions indicated in the combinatorial search. (**b**) The GA solution set from (**a**) compared to the pareto front from the combinatorial search (the solution set overlaps with the pareto front, so it is not visible on the plot other than two solutions that do not lie on the pareto front).

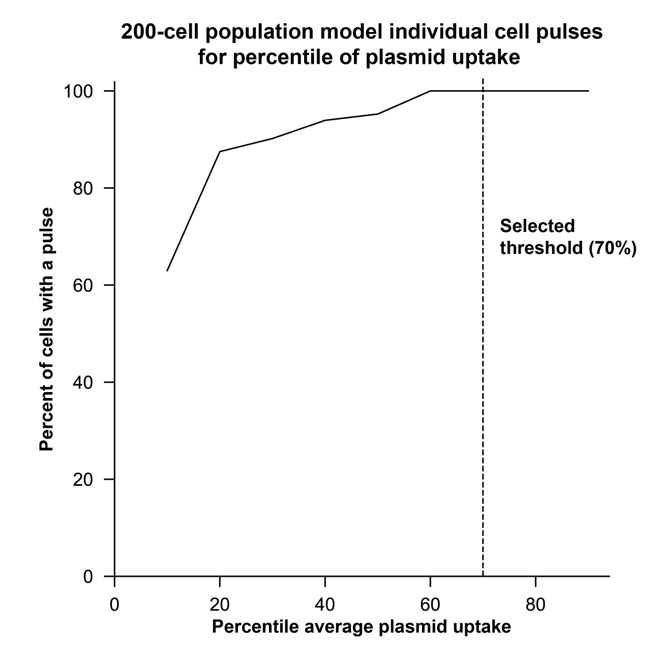

**Supplementary Figure 13. 200-cell population model individual cell pulses for percentile of plasmid uptake.** The average plasmid uptake across the 4 plasmids in the Pulse 1 experimental topology (**Figure 4b**) was calculated for each cell in the 200-cell population. Cells were defined as having a pulse if the individual cell time series had prom_rel_ > 0. For percentiles of average plasmid uptake ranging from 10^th^ percentile to 90^th^ percentile of the maximum average uptake, the percent of cells with a pulse was calculated. A threshold was selected (dashed line) to filter individual cells by plasmid uptake, such that all cells in the filtered subpopulation are hypothesized to produce a pulse.

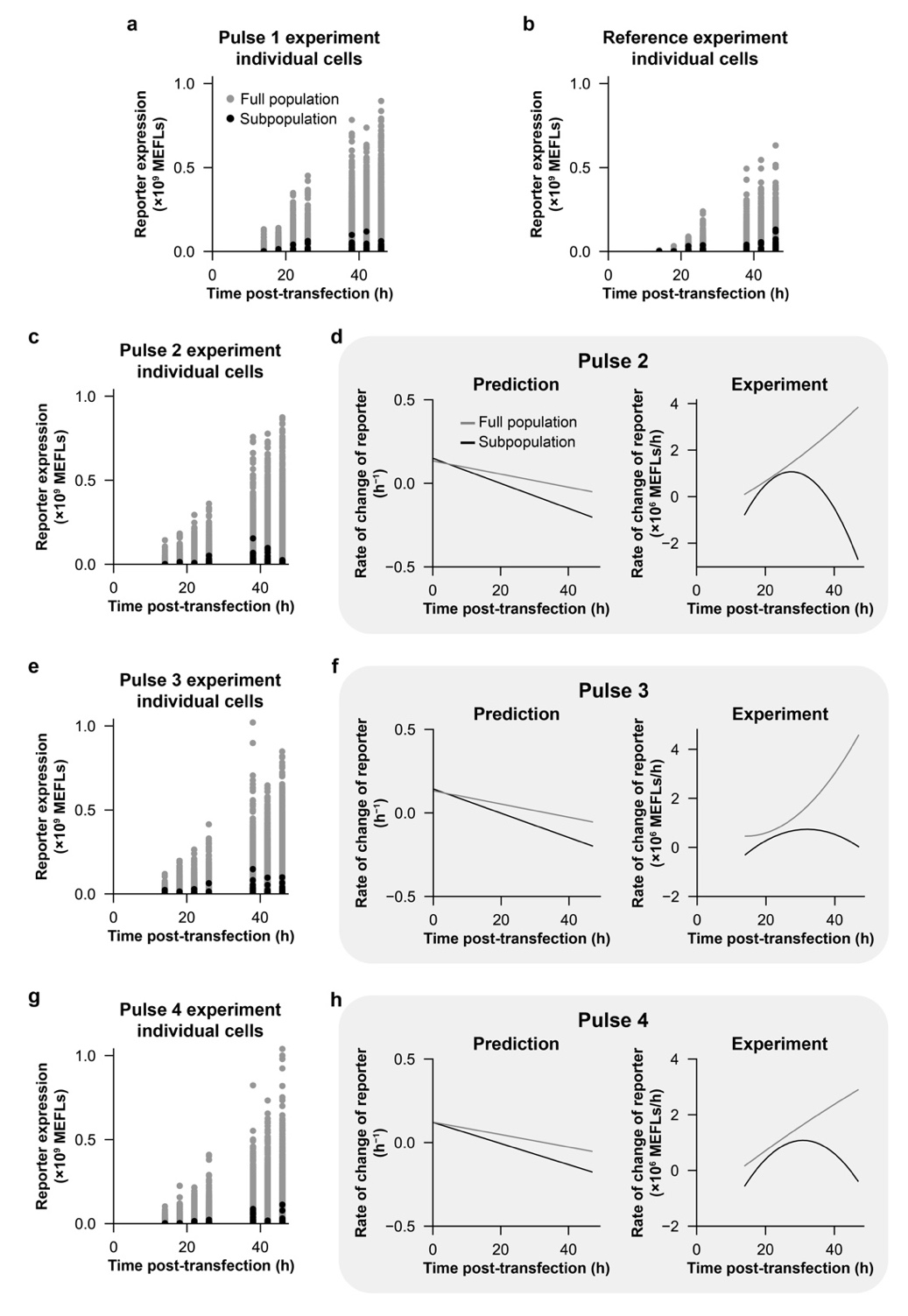

**Supplementary Figure 14. Comparison of predicted and experimentally measured Pulse Generator circuit performance.** (**a-h**) Experimental characterization of four candidate circuits plus the reference circuit and comparison to predictions. Primary experimental data are shown in panels **a**, **b**, **c**, **e**, and **g**; in each case, each dot is an individual cell within the population quantified by flow cytometry. For circuits Pulse 2–4, in panels **d**, **f**, and **h**, data were analyzed as reported and described for Pulse 1 in main **Figure 4g**. To summarize the behavior of ensembles of single cell traces (200-cell model) and focus on signatures of pulse behavior, we fit a polynomial to the reporter levels for all cells and calculated the instantaneous rate of change (first time derivative) from the polynomial fit (left). This analysis was performed for both the full population of cells and for a subset predicted to exhibit a pulse. For experimental observations, measured reporter levels were processed, both for the full population and for a subset identified by high plasmid uptake (top 30%, based on a transfection control plasmid; right).

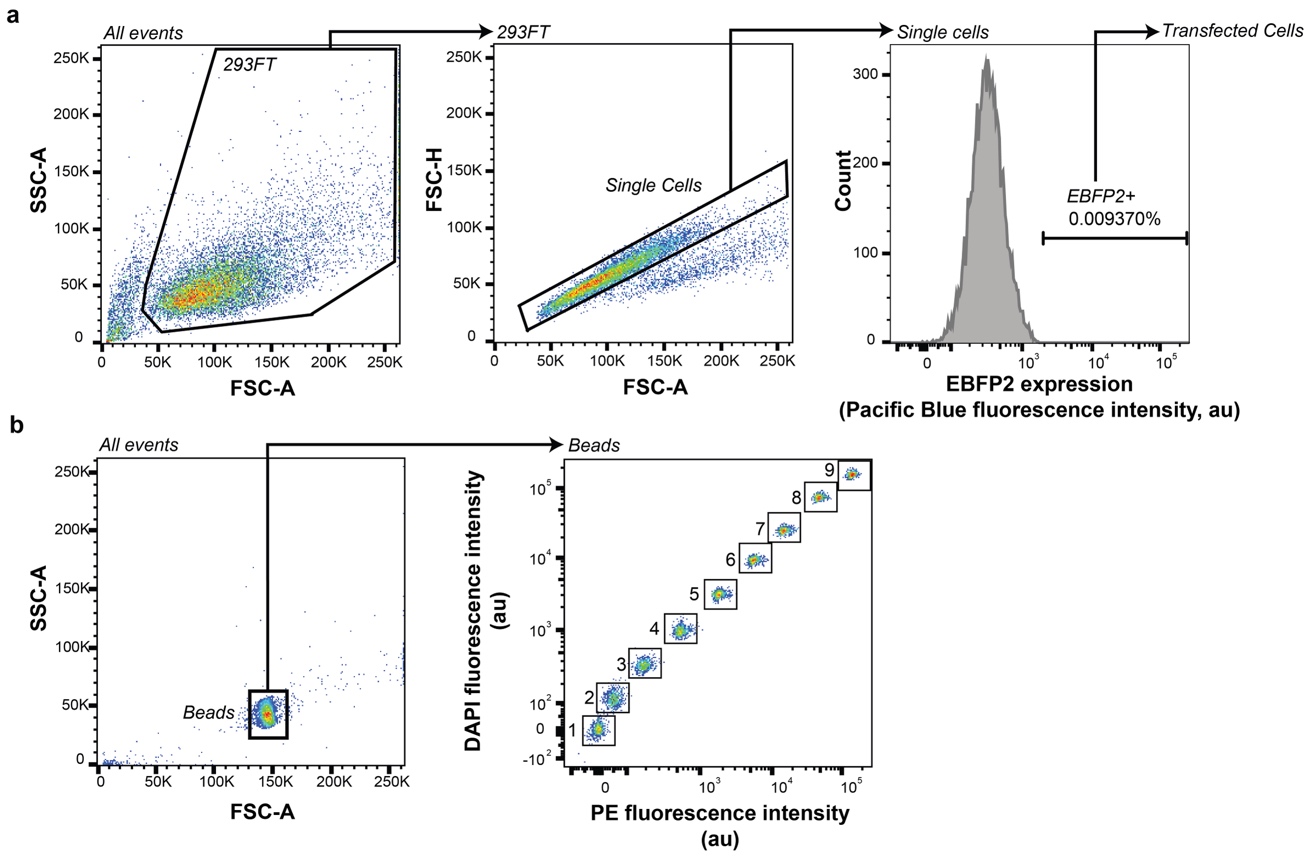

**Supplementary Figure 15. Flow cytometry gating for identifying single cells, transfected cells, and calibration beads.** (**a**) The first two plots illustrate the flow cytometry gating strategy used to identify single HEK293FT cells for a representative sample of cells. The third plot illustrates the flow cytometry gating strategy used to identify cells that are expressing EBFP2, which identifies cells that were transfected. A gate is drawn on unmodified, single HEK293FT cells to include <0.1% of cells in the Pacific Blue fluorescence channel. (**b**) Calibration of fluorescence intensities to absolute units requires inclusion of a sample of Spherotech UltraRainbow Calibration Particles (URCP) in each experiment. These beads have nine fluorescent bead populations. Beads are identified based on the FSC-A vs. SSC-A profile (left). For each experiment, two fluorescent channels were used to identify all nine bead populations (right). The mean fluorescence intensities (MFIs) of each population (gated 1-9) in the relevant channel(s) are exported and plotted against manufacturer-provided absolute values of fluorophores per bead for each population (for example, Molecules of Equivalent Fluorescein MEFLs for EYFP). To generate the calibration curve, a linear regression was performed with the constraint that the y-intercept equals zero. This calibration is done for each experiment (or time point, in the case of the pulse generator experiment), and then exported MFI values (which have arbitrary fluorescence units) are converted to absolute units using the multiplier obtained from the regression. Error from the linear regression is appropriately propagated.

**Supplementary Notes**

**Supplementary Note 1. Methods for graph-based genetic algorithms versus our method**

Graph-based GA has been proven to be useful, if not state-of-the-art, in the field of automatic design of chemical molecules. One of the earliest works on graph-based encoding is Globus et al.*^1^*. Globus et al. develop a method for drug discovery that searches for molecules similar to a target drug. A graph node (or vertex) represents an atom, and an edge represents a chemical bond type. Crossover is the only operator used to explore the search space and involves splitting parent graphs into two disconnected fragments. When a pair of new candidates are generated, they randomly replace an existing candidate in the population and enter the pool of potential parents for new candidates in the same generation. The authors find that crossover outperforms random search in finding the target molecule.

Brown et al.*^2^* further developed a comprehensive implementation of both crossover and mutation operator for graph-based GA to find novel molecular graphs. Each node can either be an atom or a predefined substructure. The method formulates two crossover operators, multipoint and subgraph. The mutation operators are divided into two categories, node mutation and edge mutation. The former category comprises four different operators, appending, pruning, inserting, and deleting. The edge mutation category comprises three operators, adding, deleting, and substituting. All three operators are implemented such that they induce the least change to the structure of the graph. The fitness function measures the similarity between a candidate with a target drug. The method utilizes multi-objective optimization to find novel molecules that are close to two target drugs. The authors demonstrate that the method is able to generate a solution set close to the Pareto front. More recent works like*^3, 4^* provide some augmentations such as mutation based on chemical reactions or application of filters before evaluating objective to remove molecules according to desired physical and chemical properties. If there are not enough candidates satisfying these filters, the process generates more molecules with mutation and crossover.

Fu et al.*^5^* proposes Reinforced GA for structure-based de novo drug design which aims to find drug molecules, or ligands, that bind tightly with a target. The method uses the outputs of neural networks to determine the probabilities of selecting parents for crossover and selecting mutation types. These neural networks are pre-trained on existing target-ligand complexes to predict binding affinity and, thus, leverage the knowledge of the shared binding physics. During the optimization process, the method uses reinforcement learning to fine-tune these networks. Similar to*^4^*, the new candidates are filtered according to properties such as solubility and toxicity before being selected to advance to the next generation.

Jensen et al.*^6^* compare the performance of graph-based GA developed in *^2, 3^* and machine learning (ML) methods such as recurrent neural network, continuous variational autoencoder, and grammar variational autoencoder for designing chemical molecules. The authors find that ML-based approaches do not outperform GA under the constraint of synthetic accessibility. Instead, the latter can obtain a better optimized objective and is computationally more efficient. Evidently, graph-based GA proves to be a favorable method for automatic design and adapting graph-based encoding to genetic circuits can be beneficial.

Our method differs from those in the chemical engineering field in two key aspects. First, chemical compounds are undirected graphs, whereas genetic circuits are directed graphs. The connectivity between two nodes in a chemical molecule characterizes a single bond type. Meanwhile, for genetic circuits, the nodes are regulatory components and there can be two different connections between any two nodes. There is also a possibility of self-regulation, that is, loop edges in the graph. As a result, using directed graph representation requires additional attention. Second, our method tackles representational redundancy in our formulation. Both graph and string-based encoding can lead to invalid circuits or multiple circuits having the same objective values. There is evidence that the encoding scheme should suppress redundant forms*^7^*. Hence, we formulate a repair operation to fix solutions with redundancy after crossover and mutation. It is possible to mimic*^3-5^* by filtering unwanted topologies and generating new ones if there are not enough candidates. However, we may be stuck in a loop and spend too much time on this step for each generation if representational redundancy is severe.

**Supplementary Note 2. Analysis of the crossover and mutation operators**

We investigated the performance of the crossover and mutation operators in GCAD to understand how the GA procedure navigates the search space and ensure the exploitative and explorative capabilities. We experimented with the Amplifier test case and ran GCAD for 120 generations with GA population size and initial population ratio being 200 and 0.25. We varied the crossover probability (p_c) and mutation probability (p_m) and for each combination of crossover and mutation probabilities, we implemented 10 independent runs (each with a different initialization seed) to capture the effect of randomization. To understand how the two operators contribute to GCAD’s search capabilities and complement one another, we varied the probability of crossover while keeping that of mutation at either 0 or 1, and vice versa.

We first inspected the diversity of the GA population in each generation, as measured by the number of unique topologies, and calculated the average percentage across generations (**Supplementary Figure 16**). When there is no mutation, the population diversity increases as the probability of crossover increases, confirming the explorative capability of the operator. However, this effect is much weaker than that of mutation, as observed in the cases where we varied the mutation probability, regardless of crossover. When the mutation probability is kept at 1, the exploitative power of crossover becomes evident as the population diversity is inversely proportional to the crossover probability.

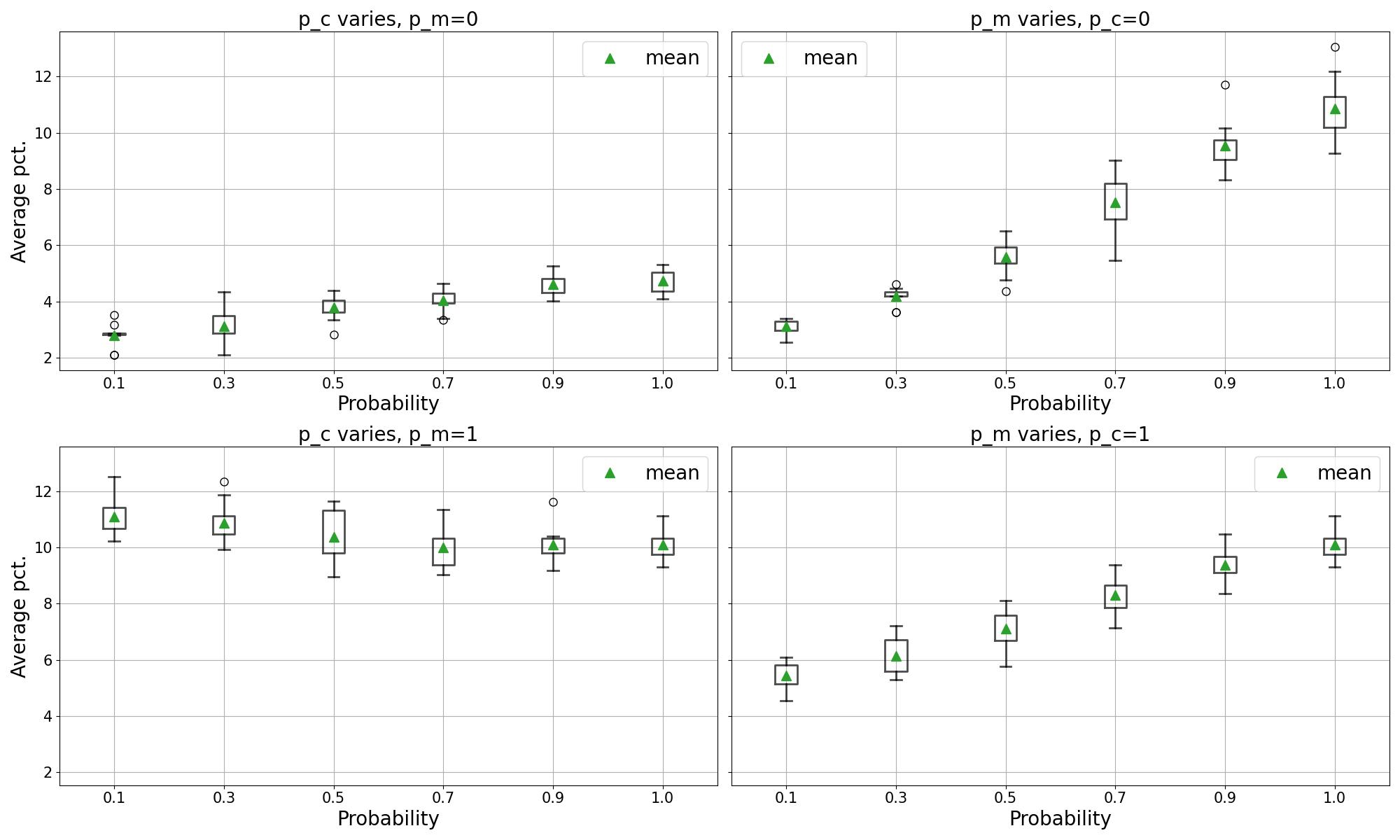

**Supplementary Figure 16. Percentage of unique topologies in the GA population averaged across 120 generations.** The distribution is over 10 different initialization seeds. Top left: the probability of crossover varies while there is no mutation. Top right: the probability of mutation varies while there is no crossover. Bottom left: We vary the crossover probability while keeping the mutation probability at 1. Bottom right: We vary the crossover probability while keeping the crossover probability at 1.

Next, we analyzed the convergence rate of GCAD for different combinations of crossover and mutation probabilities (**Supplementary Figure 17**). GCAD run is considered to converge when the number of unique topologies in the population at a generation is 1. The exploitative power of crossover is evident as GCAD converges quickly when the crossover operator is used in the GA procedure. Meanwhile, when there is only mutation, the performance is akin to a random search and takes more generations to converge.

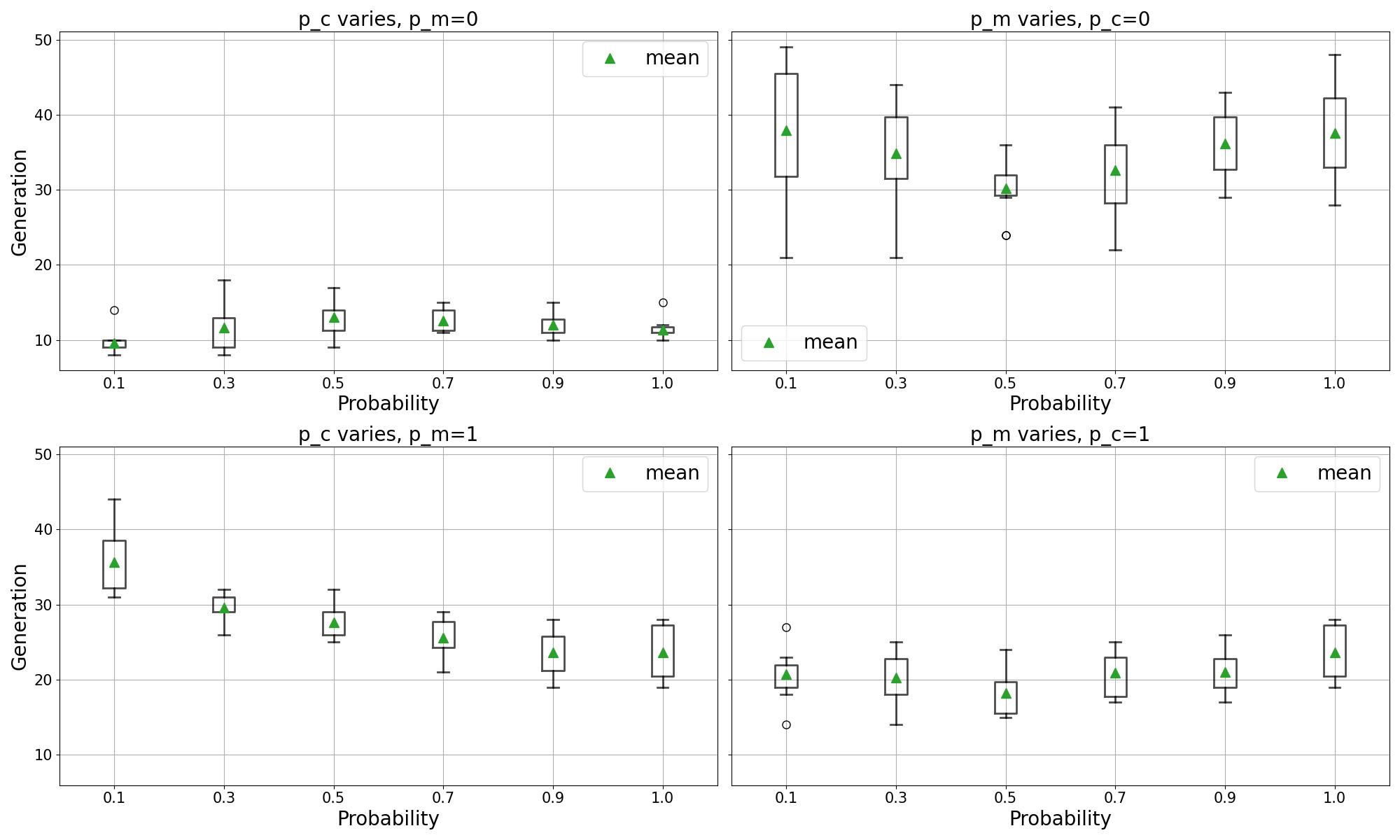

**Supplementary Figure 17. Generation at which the GA procedure converges for various combinations of crossover and mutation probabilities.** The distribution is over 10 different initialization seeds.

Finally, we evaluated how varying the strength of crossover and mutation affected the solution quality by recording the first generation in which the true optimal topology appeared in the GA population (**Supplementary Figure 18**). It is clear that even though crossover is also able to explore the search space, it is not sufficient to make meaningful search without mutation and the premature convergence results in GCAD not finding the true optimal topology. Meanwhile, the mutation operator allows us to effectively explore the search space but without crossover, the performance is unpredictable. This underscores how crossover and mutation complement each other and the need to include both operators in the GA procedure.

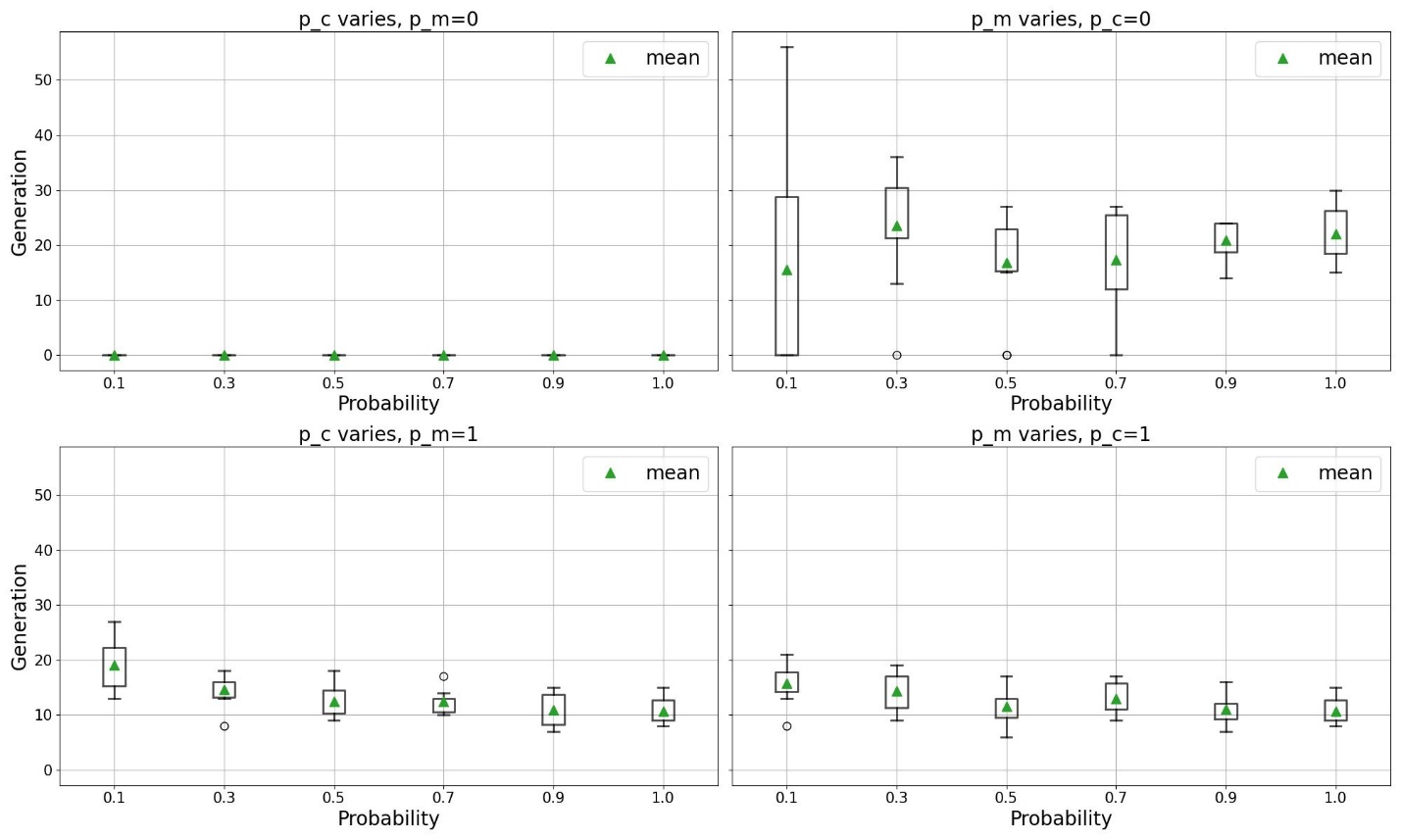

**Supplementary Figure 18. Generation at which the true optimal topology first appears in the GA population for various combinations of crossover and mutation probabilities.** The distribution is over 10 different initialization seeds.

**Supplementary Note 3. Population model GA for the Signal Conditioner test case**

We performed model selection with the 20-cell population model after the verification that the GA with the single cell model could successfully converge on a solution set with high coverage of the pareto front. We first confirmed that average simulated ON_rel_ and FI_rel_ values across a cell population for 200-cell model and the 20-cell model for a set of circuits from the single cell GA search space were consistent (**Supplementary Figure 10**). We evaluated the most optimal solutions found with the single cell GA search, circuits with mid-range FI_rel_, and circuits with low FI_rel_ to compare the population level average behavior between the two models for circuits with a diverse set of performance metrics. Since the single cell model combinatorial search and GA indicated that the best possible circuits enabled only a slight amplification in ON state and fold induction, we anticipated that this would also be the case with the population model. We ran the GA with 10 different seeds using the hyperparameters optimized for the GA with the single cell model, which revealed a similar solution set and topologies to those of the single cell model—the solution set contained a few circuits with FI_rel_ slightly greater than 1, but each of these solutions had an ON_rel_ value near 1 (**Supplementary Figure 11**). Considering this result, we chose not to further explore the full GA search space to define a confidence interval for the population model predictions, since the range of indistinguishable solutions would not provide solutions that better satisfy the design goal.

**Supplementary Tables**

**Supplementary Table 1. Plasmids obtained from external sources**

| Plasmid Name | Source | Reference |
| --- | --- | --- |
| pcDNA3.1/HygroR(+) | Thermo Fisher | V87020 |
| pEBFP2-Nuc | Addgene | RRID: Addgene_14893 |
| pLVX-TRE3G | Clontech | Takara: 631187 |
| pLVX-TetON3G | Clontech | Takara: 631187 |

**Supplementary Table 2. Plasmids used in this study, previously published by the authors**

| Plasmid# | Plasmid name | Backbone | Leonard Lab Plasmid ID | Reference |
| --- | --- | --- | --- | --- |
| pPD005 | pcDNA Golden Gate | pcDNA3.1/HygroR(+) | L1267 | RRID: Addgene_138749 |
| pPD436 | pBI-MCS-EYFP | pPD005 | L2881 | RRID: Addgene_ 58855 |
| pPD100 | ZFa43 | pPD005 | L1298 | RRID: Addgene_138761 |
| pPD270 | pZF1/ZF2 EYFP | pPD005 | L1442 | RRID: Addgene_138934 |
| pPD540 | EYFP COMET Reporter Template | pPD005 | L1270 | RRID: Addgene_138750 |
| pPD546 | pZF9 EYFP | pPD540 | L1278 | RRID: Addgene_138860 |
| pPD547 | ZF1x6c EYFP | pPD540 | L1279 | RRID: Addgene_138861 |
| pPD555 | pZF6 EYFP | pPD540 | L1287 | RRID: Addgene_138869 |

**Supplementary Table 3. Plasmids generated in this study**

| Plasmid# | Plasmid name and description | Backbone | Leonard Lab Plasmid ID | Reference |
| --- | --- | --- | --- | --- |
| **Reference Circuit** | | | | |
| **pHIE806** | pTREGV EYFP | pPD005 | L4419 | Addgene pending |
| **pHIE771** | pCMV TetON3G | pPD005 | L4386 | Addgene pending |
| **Amplifier** | | | | |
| **pHIE830** | pTREGV ZF2a | pPD005 | L4443 | Addgene pending |
| **pHIE837** | pTREGV ZF9a | pPD005 | L4450 | Addgene pending |
| **pGGB301** | pZF9/ZF6x6c ZF9a | pPD540 | L5495 | Addgene pending |
| **pGGB302** | pZF9/ZF6x6c ZF6a | pPD540 | L5496 | Addgene pending |
| **pGGB303** | pZF2x6c ZF2a | pPD540 | L5497 | Addgene pending |
| **pGGB304** | pZF2/ZF6x6c ZF6a | pPD540 | L5498 | Addgene pending |
| **pGGB305** | pZF2/ZF9x6c ZF9a | pPD540 | L5499 | Addgene pending |
| **pGGB306** | pZF9 ZF9a | pPD540 | L5500 | Addgene pending |
| **Pulse** | | | | |
| **pLR001** | pTREGV ZF1a | pPD005 | L5501 | Addgene pending |
| **pLR002** | pZF1 ZF2i | pPD005 | L5502 | Addgene pending |
| **pLR003** | pZF1/ZF2 ZF1a | pP005 | L5503 | Addgene pending |

**Supplementary Table 4. Filters for analytical flow cytometry**

| Instrument | Fluorescent proteins | Parameter/  Channel name | Excitation laser | Filter set |
| --- | --- | --- | --- | --- |
| BD LSRFortessa | EBFP2 | Pacific Blue | Violet, 405 nm | 450/50 |
|  | EYFP | FITC | Blue, 488 nm | 505LP, 530/30 |
